# Regulating *PCCA* gene expression by modulation of pseudoexon splicing patterns to rescue enzyme activity in propionic acidemia

**DOI:** 10.1101/2023.07.05.547835

**Authors:** Ulrika Simone Spangsberg Petersen, Maja Dembic, Ainhoa Martínez-Pizarro, Eva Richard, Lise Lolle Holm, Jesper Foged Havelund, Thomas Koed Doktor, Martin Røssel Larsen, Nils J. Færgeman, Lourdes Ruiz Desviat, Brage Storstein Andresen

## Abstract

Pseudoexons are nonfunctional intronic sequences that can be activated by deep intronic sequence variation. Activation increases pseudoexon inclusion in mRNA and interferes with normal gene expression. The *PCCA* c.1285-1416A>G variation activates a pseudoexon and causes the severe metabolic disorder, propionic acidemia, by deficiency of the propionyl-CoA carboxylase enzyme encoded by *PCCA* and *PCCB*. We characterized this pathogenic pseudoexon activation event in detail and identified hnRNP A1 to be important for normal repression. The *PCCA* c.1285-1416A>G variation disrupts an hnRNP A1-binding splicing silencer and simultaneously creates a splicing enhancer. We demonstrate that blocking this region of regulation with splice-switching antisense oligonucleotides restores normal splicing and rescues enzyme activity in patient fibroblasts and in a cellular model created by CRISPR gene editing. The *PCCA* pseudoexon can be exploited as a gene-regulatory switch, as healthy tissues show relatively high levels of inclusion. By blocking inclusion of the non-activated wild type pseudoexon, we increase both PCCA and PCCB protein levels, which increases the activity of the heterododecameric enzyme. Surprisingly, we can increase enzyme activity from residual levels not only in patient fibroblasts harboring *PCCA* missense variants, but also those harboring *PCCB* missense variants. This could be a potential treatment strategy for propionic acidemia.

## Introduction

Pre-mRNA splicing is the process of joining exons through excision of intervening introns. This process is catalyzed by the spliceosome and controlled by the “splicing code”, which is a network of features that constitutes a complex regulatory environment.^1,2^ The spliceosome recognizes exons from introns through core splicing signals; the 3’ and 5’ splice sites (3’ss/5’ss), and auxiliary splicing regulatory elements (SRE) that can bind splicing regulatory proteins to enhance or silence splicing in the local pre-mRNA region.^3–5^

Sequence variation that affect the splicing code will affect the outcome of the pre-mRNA splicing process and can cause disease.^6^ Deep intronic sequence variation can activate pseudoexons, which are nonfunctional intronic sequences delimited by pairs of cryptic splice sites.^7–9^ Pseudoexon activation increases the pseudoexon inclusion level in mRNA and commonly causes loss of function from frameshift and formation of premature termination codons (PTC), which can target the mRNA transcript for degradation by nonsense-mediated mRNA decay (NMD).^7,8,10^

Propionic acidemia (MIM #606054) is an autosomal recessive metabolic disorder characterized by accumulation of toxic metabolites. Patients present with metabolic decompensation and risk developing severe long-term complications affecting different organ systems. Treatment depends on metabolic control through dietary restrictions, but prevention or treatment of the complications is limited.^11,12^ Propionic acidemia occurs by deficiency of propionyl-CoA carboxylase (PCC), which is a heterododecamer enzyme composed of six alpha and six beta subunits encoded by *PCCA* and *PCCB*. The PCC enzyme catalyzes the carboxylation of propionyl-CoA to methylmalonyl-CoA in the catabolism of valine, isoleucine, threonine, methionine, odd chain fatty acids, and cholesterol in mitochondria.^13^ Propionic acidemia is genetically heterogenous and PCC deficiency can occur by deleterious sequence variation in *PCCA* or *PCCB*, with missense variants being predominant in both genes.^14–19^

Activation of an 84 bp pseudoexon (13:100305751-100305834; GRCh38) from intron 14 of the *PCCA* gene (NM_000282) has been described in a patient with a deep intronic sequence variation; c.1285-1416A>G (13:100305776; GRCh38).^20^ Inclusion of the pseudoexon introduces an in-frame PTC, triggering NMD or leading to translation of a truncated protein. Basal inclusion levels of the *PCCA* pseudoexon have previously been observed in wild type cell lines and may represent low-level aberrant splicing that is tolerated by the cell.^21^

In this study, we investigated the mechanism of the pathogenic *PCCA* pseudoexon activation event in detail to identify efficient splice-switching antisense oligonucleotides (SSO) that can block the SREs, which are responsible for inclusion of the pseudoexon in mRNA. Normal inclusion of the non-activated wild type pseudoexon reduces functional gene expression relative to the full potential, and we demonstrate that the *PCCA* pseudoexon can be exploited as a gene-regulatory switch by SSO-mediated modulation of pseudoexon splicing patterns. Curiously, we also demonstrate how overproduction of the PCCB subunit can be exploited to increase enzyme activity above normal levels by modulating the levels of PCCA. Using our most efficient SSO, we can block inclusion of the pseudoexon to increase *PCCA* gene expression, to increase both PCCA and PCCB protein levels resulting in an increase in activity of the multimeric PCC enzyme. This is a potential treatment strategy for propionic acidemia, not only for patients with pseudoexon activation, but also for patients with missense variants that have some residual activity of the enzyme.

## Results

### The pathogenic *PCCA* pseudoexon activating variation disrupts an hnRNP A1-binding splicing silencer element and simultaneously creates a splicing enhancer element

We first investigated the mechanism of activation and regulation of pseudoexon splicing to identify potential SSO target sites that can be targeted to block inclusion of the *PCCA* pseudoexon. As opposed to targeting the pseudoexon splice sites,^20^ targeting SREs is more specific with a lower risk of off-target effects. The *PCCA* c.1285-1416A>G variation has been suggested to activate the pseudoexon by creation of an exonic splicing enhancer (ESE).^20^ However, we noted that the pseudoexon activating variation also disrupts a potential hnRNP A1 core UAG/CAG motif,^22–24^ which is supported by *in silico* predictions (Figure S1 and S2).^25–27^ This is also consistent with the observation that the pseudoexon region is covered by enhanced cross-linking and immunoprecipitation (eCLIP) reads from an hnRNP A1 targeted experiment in HepG2 cells (Figure S3).^28,29^ We therefore speculated that the disease causing variation also disrupts an hnRNP A1-binding ESS, which normally functions to suppress the pseudoexon. It will thus abolish normal repression and simultaneously allow accessibility of the created ESE, which will further stimulate recognition and splicing of the pseudoexon.

We investigated the functionality of these elements by employing mutagenesis of a *PCCA* pseudoexon splicing reporter minigene (Figure 1A). The *PCCA* c.1285-1416A>G variation (MUT) has previously been demonstrated to cause complete activation in patient fibroblasts,^20^ and it efficiently activates the pseudoexon in our minigene assay (Figure 1B). An A>T variation (MUT 2) is similarly predicted to disrupt binding of hnRNP A1 but is not predicted to create an ESE (Figure S2 and S4). Accordingly, this sequence variation only causes weak activation of the pseudoexon, suggesting that disruption of the hnRNP A1-binding ESS alone is insufficient for complete activation. An A>C variation (MUT 3), also predicted to disrupt binding of hnRNP A1, is also predicted to simultaneously create an ESE and it efficiently activates the pseudoexon, indicating that the creation of an ESE is important for complete activation. Additionally, two nearby overlapping ESEs are predicted upstream from the *PCCA* c.1285-1416A>G variation. Disruption of the nearest element (MUT 4+6) results in lower inclusion levels of the activated pseudoexon, which suggests that this ESE is required for pseudoexon recognition and splicing.

**Figure 1.**
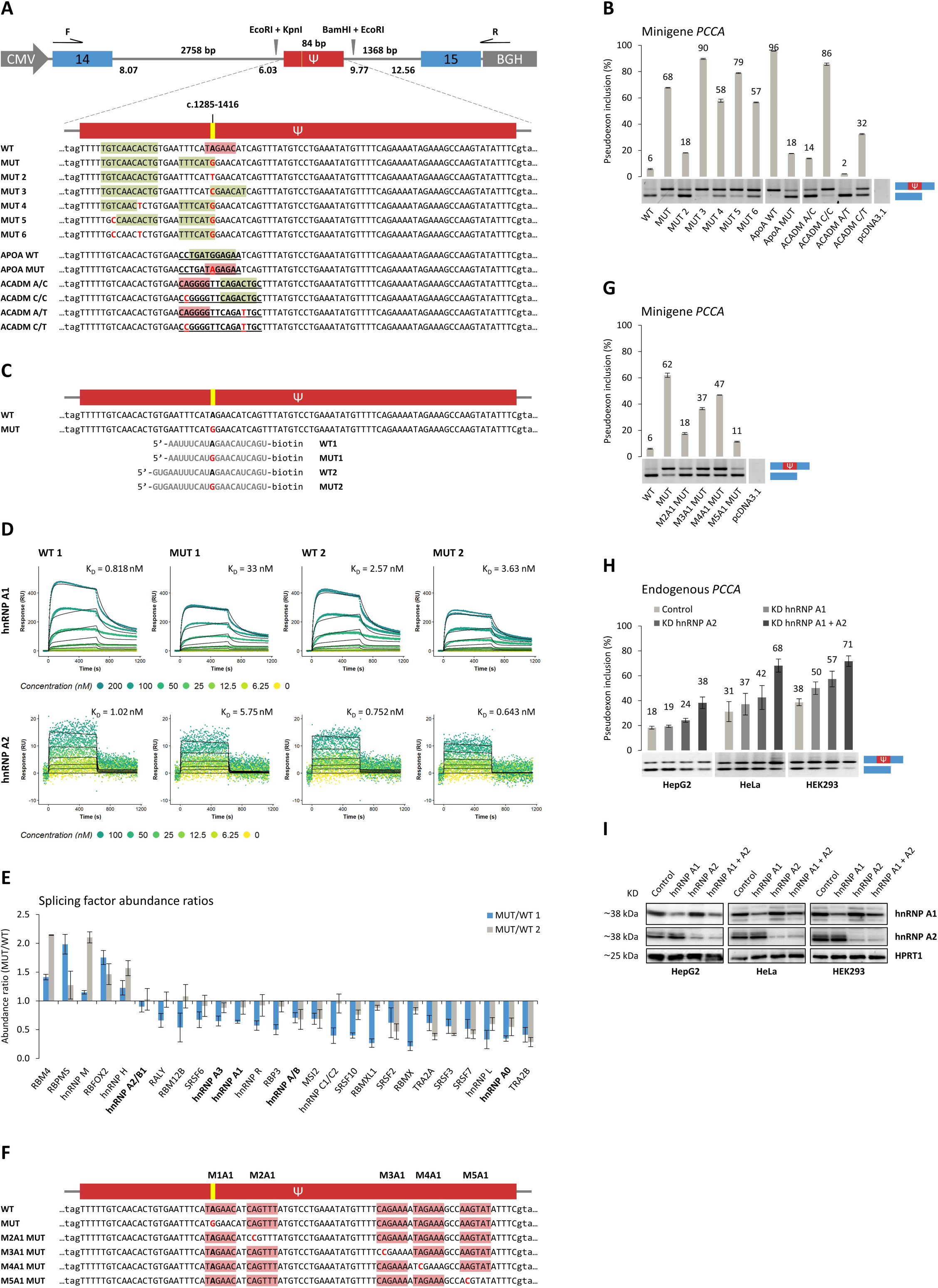
Investigating the mechanism of activation and regulation of the *PCCA* pseudoexon. **(A)** Schematic representation of the *PCCA* pseudoexon splicing reporter minigene construct and indication of mutagenesis for MUT, MUT 2-6, APOA WT/MUT, and ACADM A/C, C/C, A/T, and C/T variants of the minigene pseudoexon sequence. The minigene construct includes the *PCCA* pseudoexon, flanking introns, and neighboring exons (14+15). Location of the *PCCA* c.1285-1416A>G variation is marked in yellow. Maximum entropy (MaxEnt) scores are given for each splice site in the minigene construct. F and R indicates the location of forward and reverse primers for RT-PCR analysis of minigene *PCCA* pseudoexon splicing patterns. Exonic splicing enhancers (ESE) are indicated by green highlights, and hnRNP A1-binding exonic splicing silencers (ESS) are indicated by red highlights of the pseudoexon sequence elements. **(B)** *PCCA* pseudoexon splicing patterns from minigene variants presented in (A) transfected into HepG2 cells. Error bars indicate standard error of mean, n=3 culture wells. **(C)** Sequences of two sets of biotinylated RNA oligonucleotides covering the splicing regulatory element altered by the *PCCA* c.1285-1416A>G variation used for surface plasmon resonance imaging (SPRi) (D) and RNA affinity pulldown (E). **(D)** Binding profiles from SPRi of recombinant hnRNP A1 and hnRNP A2 with protein injections at increasing concentrations. Measurements for hnRNP A1 are fitted to a bimodal 1:2 binding model and measurements for hnRNP A2 are fitted to a monomodal 1:1 binding model. The equilibrium constant, K_D_, is given as an indicator of how well the protein binds at equilibrium. A higher K_D_ indicates that more protein is needed to reach equilibrium. The *PCCA* c.1285-1416A>G variation (MUT) results in lower binding affinity of both hnRNP A1 and hnRNP A2 relative to wild type (WT), although recombinant hnRNP A2 has a very low binding affinity to both sets of biotinylated RNA oligonucleotides and is fitted poorly to the binding model. **(E)** Top splicing factors affected by the *PCCA* c.1285-1416A>G variation (MUT) relative to wild type (WT), given by the change in abundance ratio (MUT/WT) from RNA affinity pulldown (members of the hnRNP A/B family are marked in bold). Splicing factors were included if more than two peptides were detected in all replicates and in both sets of biotinylated RNA oligonucleotides, and if p<0.05 for the change in abundance ratio in at least two replicates with applied cut-off values for the abundance ratio; <0.7 and >1.3, reached for at least one of the biotinylated RNA oligonucleotide sets. Error bars indicate standard error of mean, n=3 from individual experiments. **(F)** Indication of mutagenesis for M2A1-M5A1 MUT variants of the minigene pseudoexon sequence. Predicted hnRNP A1-binding ESSs are indicated by red highlights of the pseudoexon sequence elements. M1A1 indicates the element disrupted by the *PCCA* c.1285-1416A>G variation. **(G)** *PCCA* pseudoexon splicing patterns from minigene variants presented in (F) transfected into HepG2 cells. Error bars indicate standard error of mean, n=3 culture wells. **(H)** Endogenous *PCCA* pseudoexon splicing patterns from knockdown of hnRNP A1, hnRNP A2, and both hnRNP A1 and hnRNP A2 in HepG2, HeLa, and HEK293 cells. Error bars indicate standard deviations, n=6 culture wells from two individual experiments (HepG2), n=4 culture wells (HeLa and HEK293). **(I)** Western blots for validation of hnRNP A1 and hnRNP A2 knockdowns. CMV; cytomegalovirus promotor, Ψ; pseudoexon, BGH; bovine growth hormone polyadenylation signal, KD; knockdown.

To further support the assumption that normal repression of the *PCCA* pseudoexon depends on the presence of an hnRNPA1-binding ESS, and that creation of an ESE is required for complete activation, we replaced the SRE altered by the *PCCA* c.1285-1416A>G variation with similar previously described elements from other genomic contexts (Figure 1A and 1B). The wild type sequence from *APOA2* (NM_001643) exon 3 (APOA WT) contains an ESE that binds SRSF1 and SRSF2,^30^ and it efficiently activates the *PCCA* pseudoexon. We can disrupt this element (APOA MUT) and simultaneously create a putative hnRNP A1-binding ESS (Figure S3 and S4), illustrating the reverse change in regulation that functions to repress the pseudoexon. The wild type sequence from *ACADM* (NM_000016) exon 5 (ACADM A/C) contains an SRSF1-binding ESE and an hnRNP A1-binding ESS affected by the c.351A>C and c.362C>T variants.^31,32^ The presence of both elements causes weak activation of the *PCCA* pseudoexon, demonstrating that the creation of an ESE alone is insufficient for complete activation in the presence of an antagonizing hnRNP A1-binding ESS. When the ESS is disrupted (ACADM C/C), the pseudoexon is efficiently activated, and when the ESE is disrupted (ACADM A/T), repression of the pseudoexon is restored. When both elements are disrupted, the pseudoexon is mildly activated, again demonstrating its dependence on an ESS for normal repression. To demonstrate that the observed effects from changing these elements are general and not minigene specific, we replicated this experiment employing a different splicing reporter and obtained identical splicing patterns (Figure S5).

We then used biotinylated oligonucleotides covering the SRE affected by the *PCCA* c.1285-1416A>G variation (Figure 1C) and surface plasmon resonance imaging (SPRi) to demonstrate that the sequence variation causes a decrease in binding of hnRNP A1 (Figure 1D). We further validated this decrease in binding of hnRNP A1 (and other members of the hnRNP A/B family) by RNA affinity pulldown with subsequent analysis by mass spectrometry (Figure 1E). As indicated by the eCLIP reads covering the *PCCA* pseudoexon (Figure S3), other putative hnRNP A1-binding ESSs exist across the pseudoexon sequence (M2A1-M5A1) (Figure 1F). We investigated the functionality of these elements through inactivation by A>C mutations of the second nucleotide in the core UAG/CAG motifs, which we have previously demonstrated can abolish binding of hnRNP A1.^22,31–35^ Apart from M5A1, disruption of the putative hnRNP A1-binding ESSs are also consistent with *in silico* predictions (Figure S6). The decrease in binding of hnRNP A1 to the mutated M4A1 and M3A1+M4A1 elements was demonstrated by RNA affinity pulldown with subsequent analysis by western blotting (Figure S7). Accordingly, the pseudoexon can be mildly activated from disruption of M2A1, M3A1, or M4A1 in our minigene assay (Figure 1G). These elements likely cooperate with M1A1 in normal repression of the pseudoexon, allowing effective binding of hnRNP A1 and cross talk between the two regions of ESSs (M1A1+M2A1 and M3A1+M3A4), which is weakened when M1A1 is disrupted by the *PCCA* c.1285-1416A>G variation and antagonized by binding of serine/arginine-rich (SR) proteins to the newly created ESE that altogether causes complete activation of the pseudoexon.^36,37^ siRNA-mediated hnRNP A1 knockdown further supports dependency on hnRNP A1 in normal repression of the pseudoexon, as this results in increased inclusion levels from endogenous *PCCA* (Figure 1H and 1I). Both hnRNP A1 and hnRNP A2 are knocked down to circumvent possible effects from their compensatory relationship.^38^

### A seemingly benign single nucleotide polymorphism can activate the *PCCA* pseudoexon and may be a genetic determinant affecting *PCCA* gene expression

Because our results demonstrate that inclusion of the *PCCA* pseudoexon depends on a balance between several SREs and show that the pseudoexon can be activated from multiple sites, we investigated if single nucleotide polymorphisms (SNP) could play a role in *PCCA* expression by affecting pseudoexon inclusion (Figure 2A). Interestingly, SNP 1 increases the pseudoexon inclusion level to 21% (Figure 2B). According to Hardy-Weinberg Equilibrium, this seemingly benign SNP has an estimated carrier frequency of about 1/200 in the African/African American populations (rs186983584; C: 0.002655; gnomAD v3.1.2: https://gnomad.broadinstitute.org/), and may thus be a rare, but not negligible cause of decreased PCCA activity when found in homozygous or compound heterozygous form.

**Figure 2.**
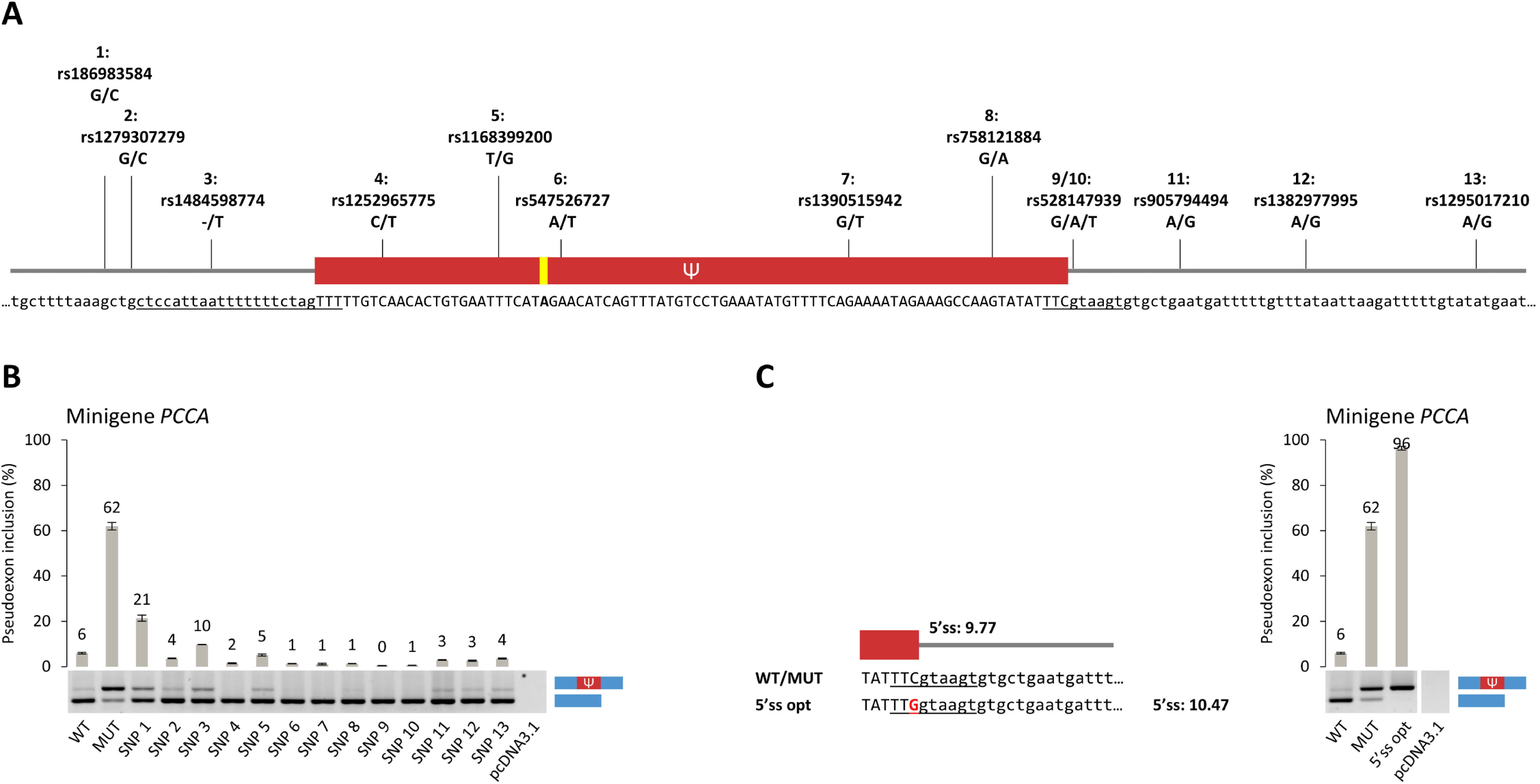
Investigating the landscape of sequence variation around the *PCCA* pseudoexon. **(A)** Schematic representation and location of single nucleotide polymorphisms (SNP) 1-13 introduced to the *PCCA* pseudoexon splicing reporter minigene. All SNPs have an alternate allele frequency <1% (gnomAD v3.1.2: https://gnomad.broadinstitute.org/). The 23-mer 3’ splice site and 9-mer 5’ splice site sequences for estimation of the maximum entropy (MaxEnt) scores are underscored. **(B)** *PCCA* pseudoexon splicing patterns from WT, MUT, and SNP 1-13 minigene variants transfected into HepG2 cells. Error bars indicate standard error of mean, n=3 culture wells. **(C)** Indication of mutagenesis for the 5’ss opt variant of the minigene pseudoexon 5’ splice site (5’ss) sequence and *PCCA* pseudoexon splicing patterns from WT, MUT, and 5’ss opt minigene variants transfected into HepG2 cells. Error bars indicate standard error of mean, n=3 culture wells. The MaxEnt score is given for the wild type and optimized 5’ss. Ψ; pseudoexon.

SNP 3 also causes low-level activation of the pseudoexon, while repression is reinforced by the presence of SNP 4 and 6-10. SNP 1 and 3 probably activates the pseudoexon by strengthening the polypyrimidine tract, and accordingly, SNP 3 increases the maximum entropy (MaxEnt) score of the 3’ss from 6.03 to 7.27. The non-activated wild type *PCCA* pseudoexon carries functional splice sites and is on the edge of increased inclusion levels.^7^ Normal repression can be abolished by changes in the balanced regulation of splicing or by small optimizations of the pseudoexon splice sites, and we demonstrate that by introducing only a small increase in the strength of the 5’ss (MaxEnt score increased from 9.77 to 10.47), we induce complete activation of the wild type pseudoexon (Figure 2C).

### Targeted SSOs can block inclusion of the *PCCA* pseudoexon to increase both PCCA and PCCB protein levels and rescue PCC enzyme activity in a cellular model

Using our *PCCA* pseudoexon splicing reporter minigene, we performed an SSO walk across the splicing regulatory region altered by the *PCCA* c.1285-1416A>G variation (Figure 3A). We identified SSOs that can restore wild type splicing patterns from the mutant minigene and block low-level pseudoexon inclusion from the wild type (Figure 3B). Consistent with our minigene results, the most efficient SSOs (SSO 1-3) also cover the upstream functional ESE, and as the SSOs can block pseudoexon inclusion from the wild type minigene, this indicates that they cover a region with overall splicing stimulatory activity, even with the presence of the hnRNP A1-binding ESS. This region was also particularly responsive when targeted by some of the 18-mer 2′-O-methoxyethyl/phosphorothioate SSOs from an extensive SSO walk across the wild type *PCCA* pseudoexon performed by Lim et al., 2020.^39^

**Figure 3.**
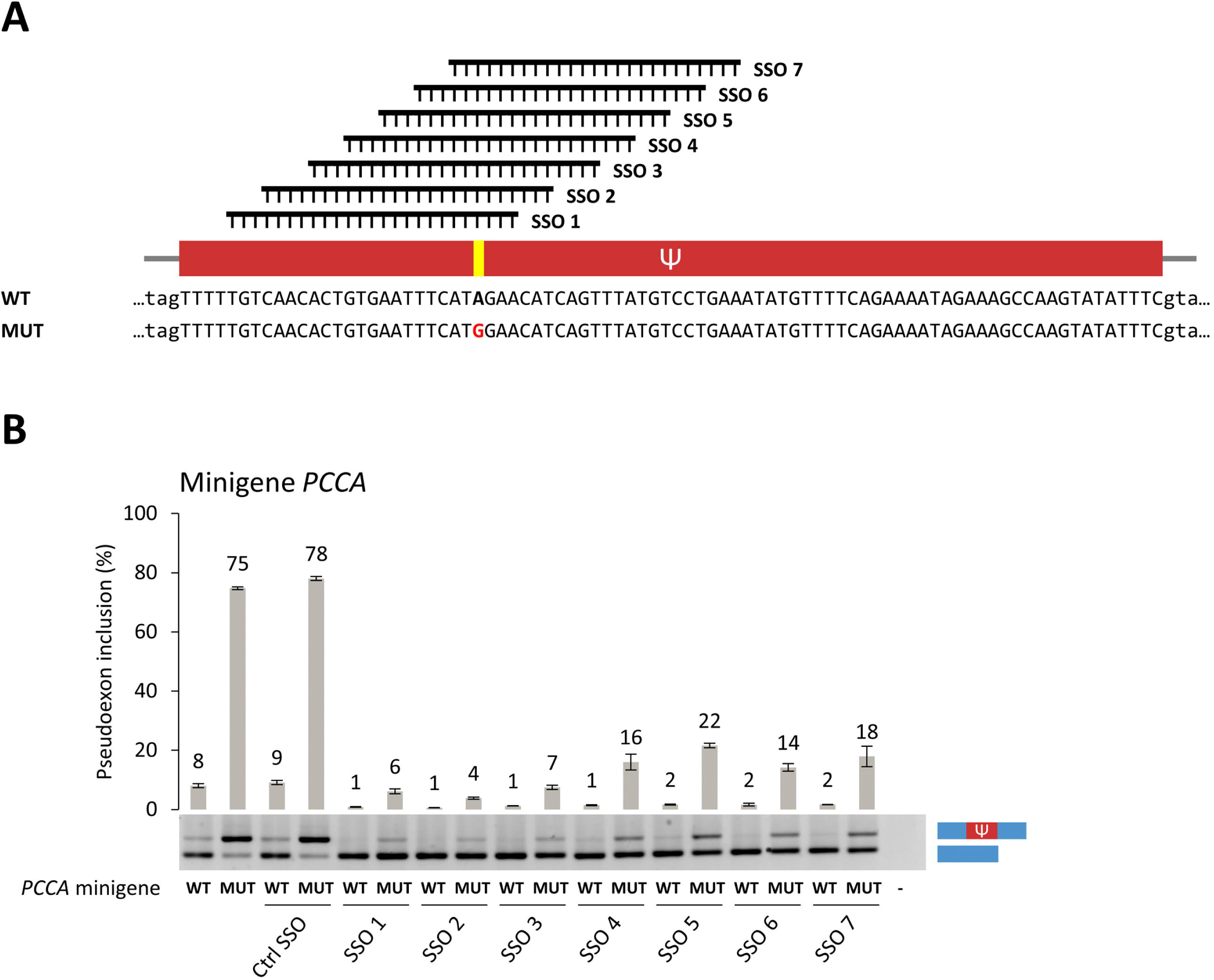
Investigating splice-switching antisense oligonucleotide (SSO)-mediated modulation of *PCCA* pseudoexon splicing patterns in minigenes. **(A)** Schematic representation of the SSO walk with location of SSO 1-7 relative to the *PCCA* pseudoexon sequence. Location of the *PCCA* c.1285-1416A>G variation is marked in yellow. **(B)** *PCCA* pseudoexon splicing patterns from WT and MUT minigene variants with additional transfection of SSO 1-7 into HepG2 cells. Error bars indicate standard error of mean, n=3 culture wells. Ψ; pseudoexon, ctrl; control.

We generated an isogenic cellular model of the *PCCA* pseudoexon activation event by using CRISPR gene editing to introduce the pathogenic pseudoexon activating variation and a 7 bp deletion of the hnRNP A1-binding ESS in HepG2 cells, which is a relevant cell line to study, as the disease manifests mainly in the liver (Figure 4A). The new HepG2 *PCCA* MUT cell line is functionally hemizygous for the *PCCA* c.1285-1416A>G variation. A large complex deletion on the other allele includes exons 13 and 14 (Figure S8), and a genomic deletion including these exons has previously been identified to cause propionic acidemia.^40^ The new HepG2 *PCCA* Del7 cell line is homozygous for a 7 bp deletion; c.1285-1411delTAGAACA, exactly covering the hnRNPA1-binding ESS.

**Figure 4.**
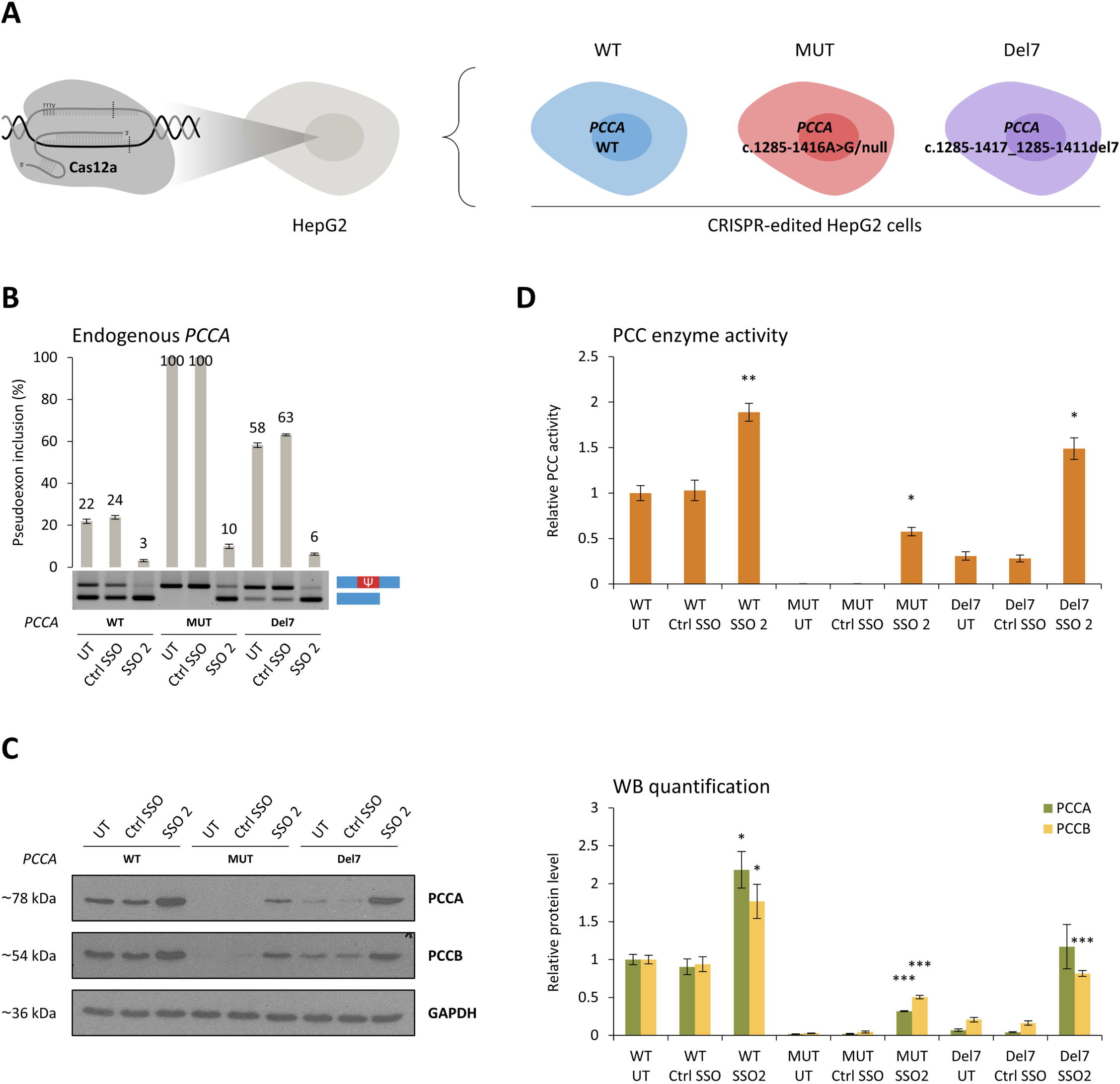
Investigating splice-switching antisense oligonucleotide (SSO)-mediated modulation of *PCCA* pseudoexon splicing patterns in a cellular model. **(A)** Schematic representation of the HepG2 *PCCA* cell lines generated with CRISPR gene editing. **(B)** Endogenous *PCCA* pseudoexon splicing patterns from untransfected (UT), control- and SSO-treated (ctrl SSO and SSO 2) HepG2 *PCCA* WT (WT), HepG2 *PCCA* MUT (MUT), and HepG2 *PCCA* Del7 (Del7) cell lines. Error bars indicate standard error of mean, n=6 culture wells from three individual experiments. SSO 2 still blocks pseudoexon inclusion efficiently in HepG2 *PCCA* Del7 even though it is only complementary to the target sequence at 18 consecutive nucleotides, having a non-complementary tail of 7 nt. **(C)** Representative western blots from investigation of PCCA and PCCB protein levels between untransfected, control-, and SSO-treated HepG2 *PCCA* cell lines, and protein quantification by densitometry normalized to the immunodetection of GADPH used as loading control, indicated as the change in protein levels relative to untransfected HepG2 *PCCA* WT. Comparisons between control- and SSO-treated samples by two-sample t-test in R; * p<0.05, ** p<0.005, *** p<0.0005. Error bars indicate standard error of mean, n=3 from individual experiments. **(D)** Propionyl-CoA carboxylase (PCC) enzyme activity between untransfected, control-, and SSO-treated HepG2 *PCCA* cell lines, indicated as the change in PCC activity relative to untransfected HepG2 *PCCA* WT. Comparisons between control- and SSO-treated samples by two-sample t-test in R; * p<0.05, ** p<0.005, *** p<0.0005. Error bars indicate standard error of mean, n=3 from individual experiments. Ctrl; control.

We performed SSO transfections of the CRISPR-edited HepG2 cells to verify the effect and efficiency by blocking inclusion of the endogenous *PCCA* pseudoexon, and to investigate the possibility to increase functional gene expression and PCC enzyme activity by targeting only the PCCA subunit. Also in this setting, the *PCCA* c.1285-1416A>G variation causes complete activation of the pseudoexon, and the splicing patterns are restored by SSO treatment (Figure 4B). The 7 bp deletion of the hnRNP A1-binding ESS efficiently activates the pseudoexon, but not to the same level of inclusion, again suggesting that disruption of this ESS is insufficient for complete activation without simultaneous creation of an ESE. Interestingly, SSO treatment causes an increase in both PCCA and PCCB protein levels in all three cell lines (Figure 4C) and consequently results in a significant increase in PCC activity (Figure 4D).

Relatively high inclusion levels of the non-activated wild type *PCCA* pseudoexon can be observed in HepG2 cells and across a human tissue panel (Figure S9). Treatment of multiple cell lines with an inhibitor of NMD, cycloheximide (CHX), verifies that some of the pseudoexon-containing transcripts are degraded (Figure S10), which implies that the pseudoexon is included at even higher levels than initially observed. Normal inclusion of this nonfunctional pseudoexon suggests that SSO-mediated modulation of the pseudoexon splicing patterns can be employed to regulate *PCCA* gene expression, as previously proposed by Lim et al., 2020.^39^ Accordingly, we demonstrate that SSO-mediated blocking of the non-activated wild type *PCCA* pseudoexon is reflected by an increase in PCCA protein levels. Remarkably, PCCB protein levels are also increased by blocking the *PCCA* pseudoexon. PCCB is synthesized in excess from stable mRNA, but the protein is unstable in the absence of PCCA.^41,42^ By increasing PCCA protein levels, excess PCCB protein is rescued, and the activity of the entire functional heteromeric PCC enzyme is thereby increased.

### SSO-mediated blocking of the *PCCA* pseudoexon can restore levels of accumulated biomarkers of propionic acidemia in a cellular model

We used RNA sequencing and differential gene expression analysis to study the transcriptomic effects of SSO treatment in our cellular model of *PCCA* pseudoexon activation in propionic acidemia. The largest difference between groups is between wild type and the two pseudoexon activating genotypes (Figure 5A), and the direction of change from SSO treatment is similar for all three cell lines. SSO treatment increases *PCCA* mRNA levels in all three cell lines, while there is no overall effect on the mRNA levels of *PCCB* (Figure 5B). By comparing log2 fold changes for genes with a significant differential expression between genotypes; HepG2 *PCCA* MUT/Del7 vs. WT, and between treatment conditions; SSO- vs. control-treated HepG2 *PCCA* MUT/Del7, we observe inverse correlations, indicating that SSO-mediated reversal of transcriptomic effects from PCCA deficiency for the two pseudoexon activating genotypes (Figure 5C).

**Figure 5.**
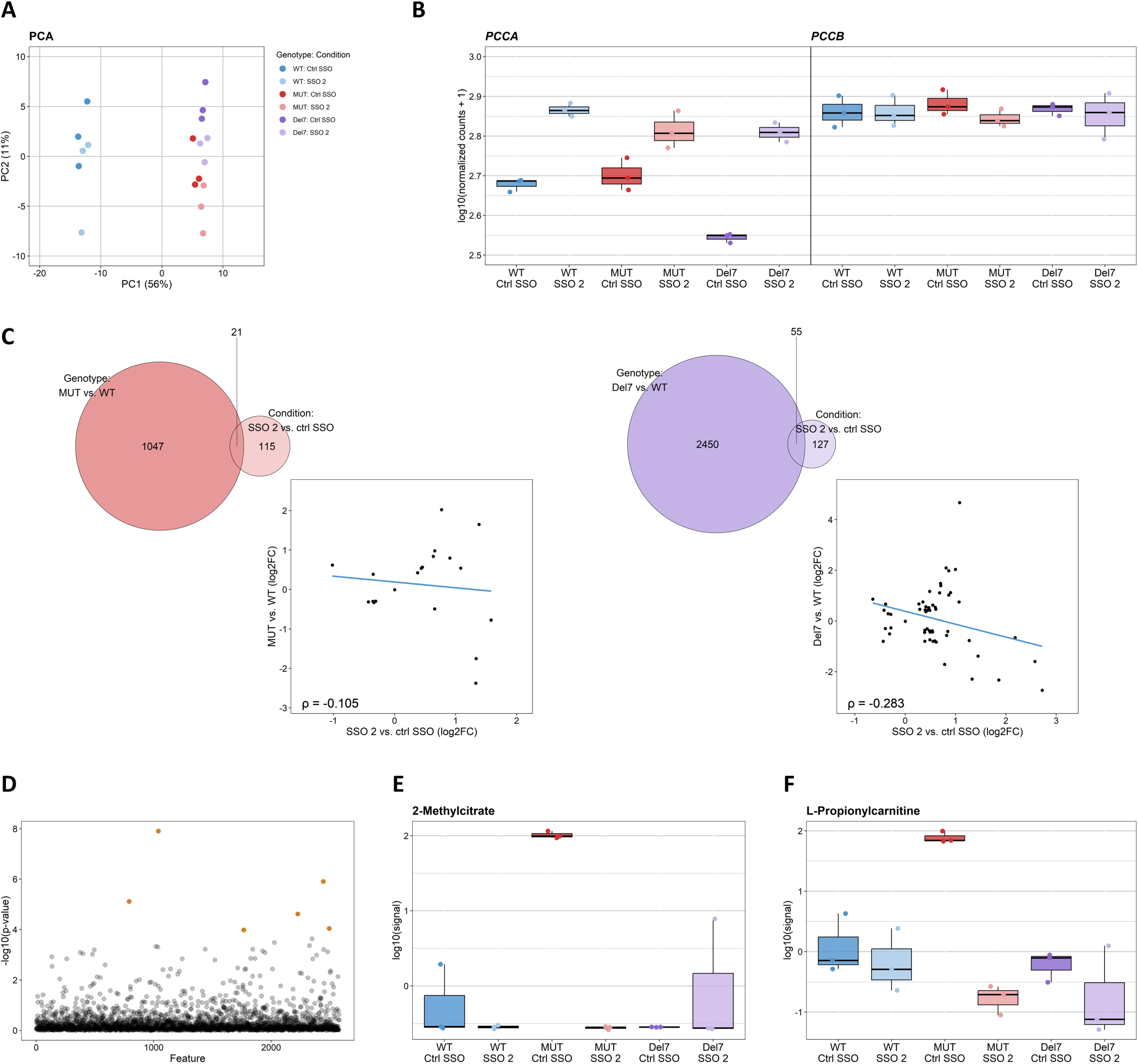
Investigating transcriptomic and metabolomic effects of the pathogenic *PCCA* pseudoexon activation event and splice-switching antisense oligonucleotide (SSO)-mediated modulation of pseudoexon splicing patterns. **(A)** Principal component analysis (PCA) plot of the variation between control- and SSO-treated (ctrl SSO and SSO 2) HepG2 *PCCA* WT (WT), HepG2 *PCCA* MUT (MUT), and HepG2 *PCCA* Del7 (Del7) cell lines in RNA sequencing data (n=3 from individual experiments). **(B)** Relative expression of *PCCA* and *PCCB* between control- and SSO-treated HepG2 *PCCA* cell lines indicated as log-transformed normalized read counts from RNA sequencing data. Adjusted p-values (p.adj.) from Wald test with post-hoc Benjamini-Hochberg test for differential gene expression between treatment conditions: p.adj.=1.742×10^−7^ (*PCCA*) and p.adj.=0.690 (*PCCB*). The expression of *PCCA* in control-treated samples is higher for HepG2 *PCCA* MUT relative to HepG2 *PCCA* Del7. Since the deletion of exon 13-14 is in-frame, this can still produce mRNA transcripts that are not targeted by nonsense-mediated mRNA decay. **(C)** Venn diagrams with overlap of significant differentially expressed genes (p.adj.<0.05) and scatter plots comparing the log2 fold changes (log2FC) of the overlapping genes between genotypes; control-treated HepG2 *PCCA* MUT/Del7 vs. WT, and between treatment conditions; in SSO- vs. control-treated HepG2 *PCCA* MUT/Del7. Pearson’s correlation, ρ, is indicated for the overlapping genes in both scatter plots. **(D)** Scatter plot of 2578 identified features in metabolomics data (n=3 from individual experiments), indicating features with a significant difference in abundance (orange) between control- and SSO-treated HepG2 *PCCA* cell lines from one-way ANOVA with post-hoc Fisher’s LSD test; p.adj.<0.05. Significant features: Neostigmine bromide (p.adj.=3.195×10^−5^), isovalerylcarnitine (p.adj.=1.609×10^−3^), pyrene-4,5-dione (p.adj.=6.627×10^−3^), 2-methylcitrate (p.adj.=0.016), L-propionylcarnitine (p.adj.=0.045), and N-3-hydroxyoctanoyl-L-homoserine lactone (p.adj.=0.045). PCA plot of the variation between control- and SSO-treated HepG2 *PCCA* cell lines in Figure S14. **(E)** Relative difference in abundance of 2-methylcitrate between control- and SSO-treated HepG2 *PCCA* cell lines indicated as log-transformed signals from metabolomics data. **(F)** Relative difference in abundance of L-propionylcarnitine. Ctrl; control.

We investigated the effect of the SSO treatment on possible off-targets from the control and SSO-treated CRISPR-edited HepG2 cells (Figure S11). 65 possible off-targets were identified with NCBI Nucleotide BLAST (https://blast.ncbi.nlm.nih.gov/Blast.cgi) of the SSO target sequence (Table S1), from which only one gene (*MIPOL1*) is differentially expressed between treatment conditions. However, no changes in the splicing patterns can be identified as affected by the possible off-target (Figure S12), and the effect on gene expression may thus be secondary to the targeted effect from the SSO. No changes in splicing patterns can be observed from other possible off-targets located within or near exons (Figure S13).

Again, using our cellular model, we also investigated the effect of SSO treatment on metabolite profiles. We identify a total of six features with a significant difference when comparing the control- and SSO-treated CRISPR-edited HepG2 cells (Figure 5D). The levels of methylcitrate and propionylcarnitine are elevated in PCCA-deficient cells with the pathogenic pseudoexon activating variation (Figure 5E). These metabolites are also accumulated in propionic acidemia,^43^ which validates the HepG2 *PCCA* MUT cell line as a cellular model of the disease. Our data show that the accumulated levels of both methylcitrate and propionylcarnitine can be restored by SSO treatment. As expected, the 7 bp deletion does not activate pseudoexon inclusion sufficiently to disrupt protein levels and enzyme activity to a degree that causes elevated levels or accumulation of these biomarkers. The HepG2 *PCCA* Del7 cell line still possesses about 30% PCC activity relative to wild type, while no enzyme activity can be measured in HepG2 *PCCA* MUT without SSO treatment.

### SSO-mediated blocking of the *PCCA* pseudoexon can increase activity of the heteromeric PCC enzyme in patient fibroblasts with missense variants

To further investigate the therapeutic potential of our most efficient SSO, we investigated the effect on PCC enzyme activity from SSO transfection into control and patient fibroblasts homozygous for the *PCCA* c.1285-1416A>G variation (Figure 6A and 6B), and into patient fibroblasts with *PCCA* or *PCCB* missense variants. Blocking of the activated *PCCA* pseudoexon results in a significant increase of PCC enzyme activity, reaching above the activity level of control fibroblasts (Figure 6D). Consistent with our observation from wild type HepG2 cells, blocking of the non-activated wild type pseudoexon also increases PCC activity in control fibroblasts. This suggests that SSO-mediated blocking of the *PCCA* pseudoexon can be employed as a potential treatment strategy for other PCCA-deficient patients with missense variants that possess some residual enzyme activity, like *PCCA* (NM_000282) c.223G>C (p.A75P), c.229C>T (p.R77W), c.412G>A (p.A138T), and c.491T>C (p.I164T).^14,44^ SSO transfection into patient fibroblasts homozygous for *PCCA* c.412G>A (p.A138T) and compound heterozygous for *PCCA* c.229C>T (p.R77W)/c.1846-2_1852del9 demonstrates a mean 4.5-fold and a mean 10-fold increase in relative PCC activity, respectively, although not significant (Figure 6D).

**Figure 6.**
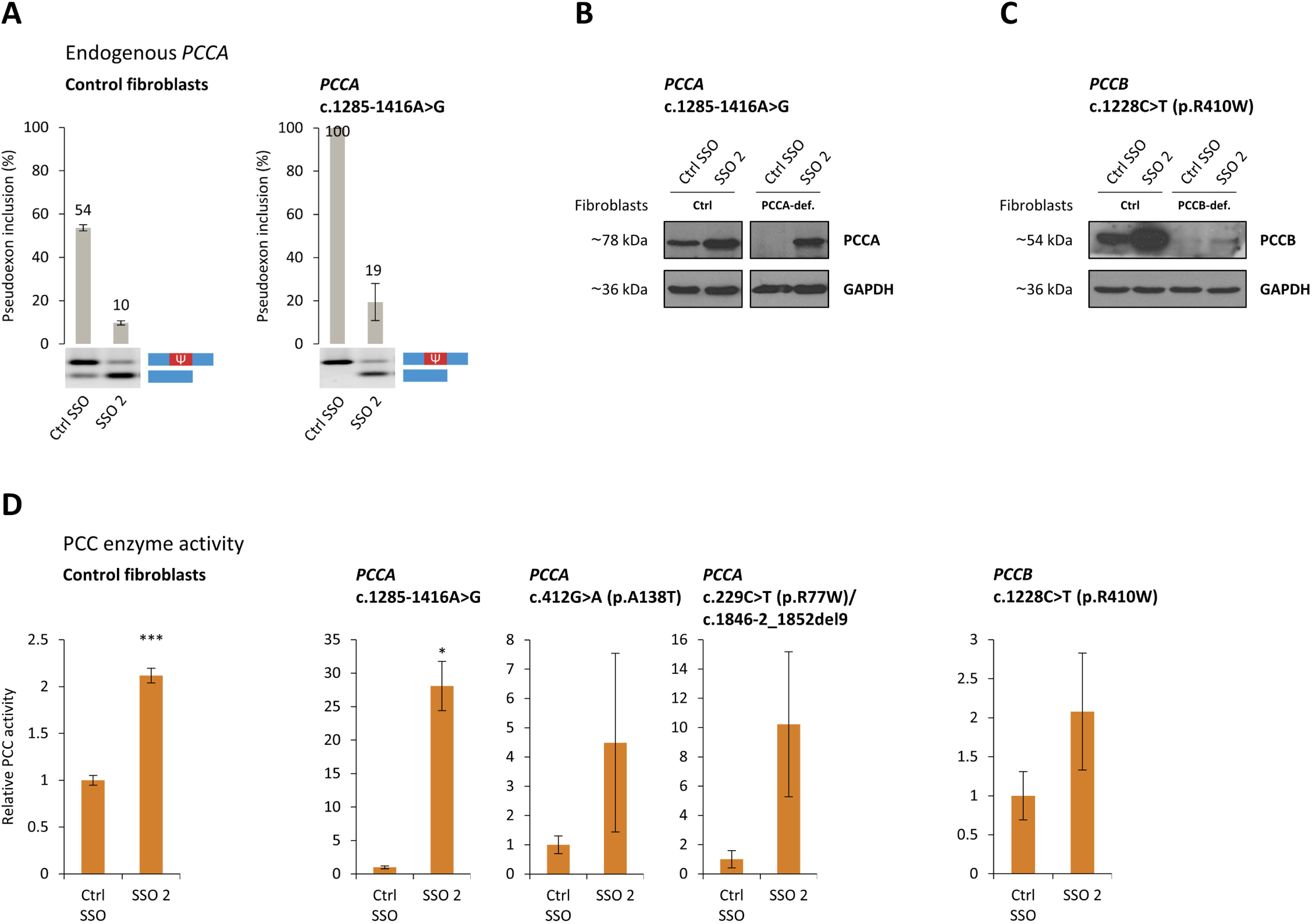
Investigating the therapeutic potential of splice-switching antisense oligonucleotide (SSO)-mediated modulation of *PCCA* pseudoexon splicing patterns. **(A)** Endogenous *PCCA* pseudoexon splicing patterns from control- and SSO-treated (ctrl SSO and SSO 2) control fibroblasts and patient fibroblasts homozygous for *PCCA* c.1285-1416A>G. Error bars indicate range, n=2 culture wells. **(B)** Representative western blots from investigation of PCCA protein levels between control- and SSO-treated control fibroblasts and patient fibroblasts homozygous for *PCCA* c.1285-1416A>G (PCCA-def.). **(C)** Representative western blots from investigation of PCCB protein levels between control- and SSO-treated control fibroblasts and patient fibroblasts homozygous for *PCCB* c.1228C>T (PCCB-def.). **(D)** Propionyl-CoA carboxylase (PCC) enzyme activity between control- and SSO-treated control fibroblasts, PCCA deficient patient fibroblasts homozygous for *PCCA* c.1285-1416A>G, homozygous for *PCCA* c.412G>A (p.A138T), and compound heterozygous for *PCCA* c.229C>T (p.R77W)/c.1846-2_1852del9, and PCCB-deficient patient fibroblasts homozygous for *PCCB* c.1228C>T (p.R410W), indicated as the change in PCC activity relative to control-treated samples. Comparisons by two-sample t-test in R; * p<0.05, ** p<0.005, *** p<0.0005. Error bars indicate standard error of mean, n=15, 4, 6, 5, and 4, respectively. The relative mean PCC activity is increased from 5% to above 100% in patient fibroblasts with *PCCA* c.1285-1416A>G compared to control fibroblasts estimated from samples from the same days of measurements. The relative mean PCC activity is increased from 3% to 5% and from 1% to 4% in patient fibroblasts with *PCCA* c.412G>A and *PCCA* c.229C>T/c.1846-2_1852del9, and from 4% to 8% in patient fibroblasts with *PCCB* c.1228C>T. Ψ; pseudoexon, ctrl; control.

Moreover, as an increase in PCCA protein levels can stabilize excess PCCB, this further suggests that SSO-mediated blocking of the *PCCA* pseudoexon can be employed as a potential treatment strategy also for PCCB-deficient patients with missense variants that possess some residual enzyme activity, like *PCCB* (NM_000532) c.1228C>T (p.R410W) and c.1606A>G (p.N536D).^15,45^ SSO transfection into patient fibroblasts homozygous for *PCCB* c.1228C>T (p.R410W) demonstrates a low but detectable increase in PCCB protein levels (Figure 6C) and a mean 2-fold increase in relative PCC activity, although not significant (Figure 6D).

## Discussion

The pathogenic *PCCA* c.1285-1416A>G variation is an example of how even a deep intronic single nucleotide sequence variation can have dramatic effects on pre-mRNA splicing patterns. The effect of deep intronic sequence variations that activate pseudoexons by alterations to the splicing regulatory environment are especially difficult to predict and may be overlooked as a cause of disease.^7^ We show that the *PCCA* pseudoexon is activated by disruption of an hnRNP A1-binding ESS and simultaneous creation of an ESE, and this dual change explains the dramatic activation of pseudoexon inclusion. Other putative hnRNP A1-binding ESSs exist across the *PCCA* pseudoexon and may cooperate in the normal repression.^36,37^ Disruption of one ESS is not sufficient to fully activate the pseudoexon, but simultaneous creation of an ESE can antagonize the remaining repression and stimulates complete pseudoexon inclusion. We demonstrate that the balanced regulation of splicing can be shifted to activate the pseudoexon for inclusion in mRNA from multiple sites, as we have previously demonstrated with mutagenesis of another splicing reporter minigene representing the splicing patterns of an *MTRR* pseudoexon.^33^ Inclusion of the minigene *PCCA* pseudoexon is increased by introducing a seemingly benign SNP; rs186983584 (C). This SNP may be a genetic determinant of a lower-level pseudoexon activation event that will affect the functional expression of *PCCA*, like other deep intronic sequence variations have been identified as modifiers of gene expression and possible disease risk factors as a consequence of their impact on splicing regulation and the use of cryptic splice sites.^46–51^

By blocking inclusion of the *PCCA* pseudoexon using an SSO targeting the splicing regulatory region altered by the pathogenic pseudoexon activating variation, we can increase the activity of the heterododecameric PCC enzyme by modulating the levels of just one subunit, as *PCCA* expression is a limiting factor for both PCC subunits. This suggests that the activity of other multi-subunit proteins also may be increased by increasing the levels of functional mRNA of only one of the subunits, despite the apparently fixed stoichiometry of the subunits in the functional protein. It illustrates a principle, where increasing the functional mRNA amounts encoding only one subunit is matched by increased rescue of the other subunit at the protein level. We demonstrate that the PCC enzyme activity can reach the level of controls in patient fibroblasts homozygous for the *PCCA* c.1285-1416A>G variation and about 50% activity relative to wild type in the new functionally hemizygous HepG2 *PCCA* MUT cell line, and that this is sufficient to restore the levels of accumulated biomarkers of propionic acidemia. An inability to reach wild type PCC activity levels from transfection of a wild type *PCCA* expression vector into PCCA-deficient patient fibroblasts,^14,52^ or from injection of a *PCCA*-expressing adeno-associated virus (AAV) vector into the *Pcca*^−/−^ (A138T) mouse model that produces higher levels of PCCA protein relative to wild type,^53^ have previously been suggested to be due to the imbalanced expression of the two subunits.^14,53^ Accordingly, a much greater increase in PCC enzyme activity have been demonstrated from dual *PCCA* and *PCCB* mRNA transfected into patient fibroblasts at an optimal molar ratio of 1:1, compared to transfection of single mRNA.^54^ In this study, we demonstrate a significant increase in PCC activity reaching normal levels by SSO treatment in cell lines with *PCCA* pseudoexon activation, which is thereby not limited from modulation of the endogenous expression.

Splice-switching using antisense oligonucleotides has emerged as a promising new strategy in treatment of genetic diseases,^55–59^ and pseudoexons are optimal targets for SSO treatment, as generation of wild type mRNA transcript will result from blocking pseudoexon inclusion. SSO-mediated blocking of an increasing number of disease-associated pseudoexons have been demonstrated to restore enzyme activity.^20,33,60–62^ Delivery of antisense oligonucleotide molecules still poses a major challenge and is limited to a few organs, like liver, kidneys, CNS, and eyes.^63^ Ligand conjugation strategies can optimize delivery by inducing receptor-mediated uptake in target cells. Antisense oligonucleotides will accumulate in liver and kidneys from systemic administration,^64^ and the hepatocyte uptake and drug potency can be enhanced through N-acetylgalactosamine (GalNAc) conjugation targeting the asialoglycoprotein receptor (ASGPR).^65,66^

We demonstrate that the *PCCA* pseudoexon can be exploited as a gene-regulatory switch, as SSO-mediated modulation of the wild type pseudoexon splicing patterns can regulate *PCCA* gene expression. Due to the genetic heterogeneity of propionic acidemia, no clear genotype-phenotype correlations have been discovered in studied patients.^16–19^ Nevertheless, residual PCC enzyme activity appear to determine disease severity, although with notable exceptions.^16,18^ In this study, SSO-mediated blocking of the non-activated wild type *PCCA* pseudoexon results in increased activity of the heterododecameric PCC enzyme in patient fibroblasts with pathogenic *PCCA* or *PCCB* missense variants, although the overall enzyme activity is still low compared to the relative activity in control fibroblasts. The initial residual enzyme activity and the effect from SSO-mediated blocking of the *PCCA* pseudoexon may be higher in other tissues more relevant to treatment, like liver, which is the major organ of propionate metabolism and has a higher expression of the PCC subunits. Moreover, it may be possible to alleviate symptoms of propionic acidemia with just a small increase in PCC enzyme activity. Liver-specific supplementation by transgenic expression of human *PCCA* resulting in only 10-20% of wild type PCC activity was sufficient to prevent lethal ketoacidosis in early infant periods for the fatal *Pcca*^−/-^ mouse model,^67^ and we observed normalization of metabolites in our HepG2 *PCCA* Del7 cells that had 30% of control enzyme activity. SSO-mediated blocking of the non-activated wild type *PCCA* pseudoexon can potentially contribute to alleviate symptoms in both PCCA- or PCCB-deficient patients with missense or leaky splicing variants^68,69^ that allow for some residual PCC enzyme activity. This represents an example of targeted augmentation of nuclear gene output (TANGO),^39^ which has shown great potential to increase functional *SCN1A* expression by SSO-mediated blocking of a naturally occurring non-productive alternative splicing event in treatment of Dravet syndrome.^39,70,71^ It is plausible that many other pseudoexons can be exploited as gene-regulatory switches in treatment of other, not only, monogenic diseases.

## Materials and Methods

### Cell cultures

HepG2, HeLa, HEK293, and fibroblast cell lines were cultured in RPMI-1640 medium (Sigma-Aldrich; United Kingdom) supplemented with 10% fetal bovine serum, L-glutamine (4 mM), and penicillin (100 U/ml)/streptomycin (100 µg/ml) at 37°C in 5% CO_2_. We have used four skin fibroblast cell lines derived from propionic acidemia patients with different genotypes; homozygous for *PCCA* c.1285-1416A>G,^20^ homozygous for *PCCA* c.412G>A (p.A138T),^44,72^ compound heterozygous for *PCCA* c.229C>T (p.R77W)/c.1846-2_1852del9,^44,72^ and homozygous for *PCCB* c.1228C>T (p.R410W) (not previously described).

### CRISPR gene editing

New HepG2 cell lines; HepG2 *PCCA* MUT, HepG2 *PCCA* Del7, and HepG2 *PCCA* WT, were generated using the Alt-R CRISPR-Cas12a (Cpf1) system (Integrated DNA Technologies; Coralville, IA) for homology-directed repair. A CRISPR RNA (crRNA) guide sequence with high specificity for the target site was designed using the Breaking-Cas web tool^73^ (https://bioinfogp.cnb.csic.es/tools/breakingcas). crRNA (PCCA-PE.MUT.crRNA: 5’-UAAUUUCUACUCUUGUAGAUAGGACAUAAACUGAUGUUCUAUG-3’) and Cas12a ribonucleoprotein was co-transfected with a 75 nt single-stranded DNA donor template of the *PCCA* c.1285-1416A>G sequence variant (−37/+37 nt) into HepG2 cells at 10 nM and 3 nM concentrations using Lipofectamine CRISPRMAX Transfection Reagent (Invitrogen; Carlsbad, CA). Cell cultures were expanded from single-cell colonies and gene editing was confirmed by Sanger sequencing (Figure S8). The HepG2 *PCCA* MUT cell line is functionally hemizygous for the *PCCA* c.1285-1416A>G variation with a large complex deletion on the other allele, spanning from intron 12 to intron 14, including the pseudoexon. The HepG2 *PCCA* Del7 cell line is homozygous for a 7 bp deletion; c.1285-1411delTAGAACA, and the HepG2 *PCCA* WT cell line derives from a negative clone in which no *PCCA* gene editing has occurred.

### Minigene constructs

The *PCCA* pseudoexon splicing reporter minigenes contain *PCCA* exon 14, intron 14, and exon 15 between NheI and XhoI restriction sites in the pcDNA3.1(+) plasmid vector. Restriction sites; EcoRI+KpnI and BamHI+EcoRI, were introduced for easy mutagenesis by insertion of short sequences 98 bp upstream (GAATTCTGGT) and 131 bp downstream (GATCCGAATTC) of the *PCCA* pseudoexon sequence within intron 14. Mutagenesis was performed by Synbio Technologies, Monmouth Junction, NJ, and all insert sequences was confirmed by Sanger sequencing.

### SSO, siRNA and minigene transfections

Cells were grown to or seeded at a density of 60-80% confluency on the day of transfection. SSOs were transfected at the time of cell seeding using Lipofectamine RNAiMAX Transfection Reagent (Invitrogen; Carlsbad, CA) with final SSO concentrations of 20 nM. All SSOs used are RNA oligonucleotides with 2’-O-methyl modifications and phosphorothioate backbones (LGC Biosearch Technologies; Risskov, Denmark) (Table S2). A non-targeting sequence was used as SSO control. Knockdown was performed by double transfections of ON-TARGETplus SMARTpool siRNA targeting hnRNP A1 (L-008221-00; Dharmacon; Lafayette, CO) and hnRNP A2/B1 (L-011690-01; Dharmacon; Lafayette, CO), using non-targeting siRNA (D-001810-10; Dharmacon; Lafayette, CO) as control. Transfections of siRNA were performed at the time of cell seeding and again after 48 h using Lipofectamine RNAiMAX Transfection Reagent with final siRNA concentrations of 40 nM at both transfections. Plasmid minigenes were transfected 24 h after cell seeding using X-tremeGENE 9 DNA Transfection Reagent (Roche; Mannheim, Germany). HepG2, HeLa, and HEK293 cells were transfected in 12-well plates (3.5 cm^2^/well) for RNA analysis, in 6-well plates (9.6 cm^2^/well) for protein analysis, and in 10 cm dishes (56.7 cm^2^) for metabolite analysis. Fibroblasts were transfected in 6-well plates (9.6 cm^2^/well) for both RNA and protein analysis. Cells were washed in phosphate buffered saline and harvested for RNA analysis 48 h after SSO transfections, 72 h after the first siRNA transfections, and 48 h after minigene transfections with no additional SSO or siRNA transfections. Cells were harvested for protein analysis 72 h after SSO transfections and the first siRNA transfections. For cells treated with CHX, media was changed and replaced with new media containing 40 µg/ml CHX 6 h before cell harvest.

### RNA analysis by RT-PCR

Cell lysis was performed by adding TRIzol Reagent (Ambion; Carlsbad, CA). After homogenization of cell lysates, total RNA was extracted by phase separation with chloroform and precipitation in isopropanol. cDNA was synthesized from 0.5 µg RNA using the High-Capacity cDNA Reverse Transcription Kit (Applied Biosystems; Foster City, CA) in 10 µl reactions. To investigate splicing patterns of the minigene *PCCA* pseudoexon, the region from *PCCA* exon 14 to the BGH termination sequence of the pcDNA3.1(+) backbone was amplified (PCCA.ex14.F: 5’-ACCCCTACAAGTCTTTTGGTTTAC-3’ and BGH.R: 5’-AACTAGAAGGCACAGTCGAGGCTG-3’) from 1 µl cDNA using TEMPase Hot Start 2x Master Mix (Ampliqon; Odense M, Denmark) in 10 µl PCR reactions. To investigate splicing patterns of the endogenous *PCCA* pseudoexon, the region from *PCCA* exon 14 to exon 15 was amplified (PCCA.ex14.F and PCCA.ex15.R: 5’-TTCCTGGTTGGATGCCACTG-3’). PCR program: 95°C for 15 min, 30 cycles at 95°C for 30 sec, 60°C (minigene)/51°C (endogenous) for 30 sec, and 72°C for 20 sec, followed by 72°C for 5 min. PCR samples were visualized on 1.5% SeaKem LE agarose (Lonza; Rockland, ME) gels containing 0.5x GelRed (Biotium; Fremont, CA), and pseudoexon inclusion was quantified from molar ratios between PCR products using the Fragment Analyzer system (Advanced Analytical Technologies; Ames, IA).

### Protein analysis by western blotting

Cells were disrupted by freeze-thawing in lysis buffer (10 mM Tris-HCl pH 7.5, 150 mM NaCl, 0.1% Triton X-100, 10% glycerol) with protease inhibitor and centrifuged for 10 min at 4°C. The supernatant was collected, and protein concentration was determined by the Bradford method (Bio-Rad Laboratories; Hercules, CA). Equal amounts of lysed extracts were loaded on a 10% sodium dodecyl sulfate (SDS)-polyacrylamide gel. After electrophoresis, proteins were transferred to a nitrocellulose membrane (iBlot 2 Transfer Stacks, mini) in an iBlot 2 Gel Transfer Device (Invitrogen; Carlsbad, CA). Immunodetection was carried out using commercially available primary antibodies against PCCA (1:500; sc-374341; Santa Cruz Biotechnology; Santa Cruz, CA) or PCCB (1:500; sc-393929; Santa Cruz Biotechnology; Santa Cruz, CA), and GAPDH (1:5000; ab8245; Abcam; Cambridge, UK) as loading control, with a secondary antibody against mouse IgG (1:2000; #7076; Cell Signaling Technology; Danvers, MA). Antibody binding was detected by enhanced chemiluminescence (GE Healthcare; Buckinghamshire, United Kingdom). Protein quantification was performed using a GS-900 Calibrated Densitometer (Bio-Rad Laboratories; Hercules, CA).

For siRNA knockdown experiments, cell lysis was performed by adding M-PER lysis buffer (Thermo Scientific; Rockford, IL) supplemented with 1x cOmplete Protease Inhibitor Cocktail (Roche; Mannheim, Germany) and 1mM phenylmethylsulfonyl fluoride (PMSF). Protein concentration was estimated using the Pierce BCA Protein Assay Kit (Thermo Scientific; Rockford, IL). Equal amounts of lysed extracts were loaded on a NuPAGE 4-12% Bis-Tris Gel (Invitrogen; Carlsbad, CA) and transferred to a polyvinylidene difluoride (PVDF) membrane. Immunodetection was carried out using primary antibodies against hnRNP A1 (1:1000; R9778; Sigma-Aldrich; St. Louis, MO) or hnRNP A2/B1 (1:1000; sc-53531; Santa Cruz Biotechnology; Santa Cruz, CA), and HPRT1 (1:1000; HPA006360; Sigma-Aldrich; St. Louis, MO) as loading control.

### PCC enzyme activity assay

PCC activity was assayed by measuring the enzyme-dependent incorporation of [^14^C]-bicarbonate into non-volatile products from the Krebs cycle as previously described.^74^ Briefly, cells were resuspended in 20 mM Tris-HCl pH 8.0 with 0.81 mM glutathione and disrupted by freeze-thaw cycles. The enzymatic reaction was initiated with addition of the homogenate, maintained at 30°C for 20 min, and stopped with 30% trichloroacetic acid. The samples were centrifuged at 16,873 ×g for 15 min, the supernatant was transferred to a scintillation vial and subjected to an evaporation process at room temperature for 24-48 h. The non-volatile products were measured in a Tri-Carb 2810 TR Liquid Scintillation Counter (PerkinElmer; Waltham, MA) after 48 h. PCC activity is expressed in pmols of incorporated [^14^C] min^−1^ mg^−1^ of total protein.

### RNA affinity pulldown and protein quantification by mass spectrometry

Two sets of 3’-end biotinylated RNA oligonucleotides (LGC Biosearch Technologies; Risskov, Denmark) (Table S3) covering the same region of the *PCCA* pseudoexon was used to perform affinity purification of RNA-binding proteins as previously described.^32^ 1000 pmol of each 3’ end biotin-coupled RNA oligonucleotide was immobilized in 50 µl streptavidin-coupled magnetic beads; Dynabeads M-280 Streptavidin (Invitrogen; Oslo, Norway), and incubated with HeLa nuclear extract (CILBiotech; Mons, Belgium). Tandem Mass Tag 6-plex labelling of eluted RNA-binding proteins was performed using the TMTsixplex Isobaric Mass Tagging Kit (Thermo Scientific; Rockford, IL) and analysis by liquid chromatography tandem mass spectrometry (LC-MS/MS) was performed as previously described.^75^

Or proteins were eluted in XT sample buffer (Bio-Rad Laboratories; Hercules, CA) and analyzed by western blotting with immunodetection using primary antibodies against hnRNP A1 (1:2000; R9778; Sigma-Aldrich; St. Louis, MO), hnRNP A2/B1 (1:1000; sc-53531; Santa Cruz Biotechnology; Santa Cruz, CA), TDP-43 (1:1000; 10782-2-AP; Proteintech; Rosemont, IL), and hnRNP K (1:1000; sc-28380; Santa Cruz Biotechnology; Santa Cruz, CA).

### Surface plasmon resonance imaging

SPRi was carried out as previously described.^75^ Briefly, biotinylated RNA oligonucleotides (LGC Biosearch Technologies; Risskov, Denmark) (Table S3) were immobilized onto a G-strep sensor chip (Ssens; Enschede, Netherlands) for 20 min. The following recombinant proteins were injected for 10 min, followed by dissociation for 8 min; 6.25-200 nM hnRNP A1 (TP303314; OriGene Technologies; Rockville, MD) and 6.25-100 nM hnRNP A2 (TP319318; OriGene Technologies; Rockville, MD). A continuous flow of SPR buffer (10 mM Tris-HCl pH 7.9, 150 mM KCl, 3.4 mM EDTA, 0.005% Tween-20) was injected both before, during, and after protein injections. Nuclear extract was used as a control for the binding efficiency of oligonucleotides. Binding hnRNP A2 was fitted to a 1:1 kinetics model with Scrubber2 (v.2.1.1.0, BioLogics). For hnRNP A1 a biphasic 1:2 model was used in ClampXP (v.3.50, Biosensor Data Analysis).

### RNA sequencing and analysis

cDNA libraries were prepared from 500 ng RNA using the NEBNext Ultra II Directional RNA Library Prep Kit (New England Biolabs; Ipswich, MA) with rRNA depletion. Libraries were quantified by qPCR using the Kapa Library Quantification Kit (Roche; Cape Town, South Africa), and the pooled library (from 27 RNA samples) was loaded to an SP flow cell and sequenced on the NovaSeq 6000 platform (Illumina; San Diego, CA) with 2 x 150 bp paired-end sequencing. RNA sequencing reads were mapped to the GRCh38 reference genome using STAR (v.2.6.1a).^76^ Differential gene expression analysis was performed with DESeq2,^77^ using read counts overlapping genes of the Ensembl (release 93)^78^ annotation reference.

### Metabolite extraction and metabolomics

Cells were washed twice in cold 10 mM ammonium acetate, scraped off the culture dish, pelleted, and frozen at −80°C before metabolite extraction. 1 ml ice-cold extraction solvent (50% methanol; 30% acetonitrile; 20% water) was added to the cell pellets. The mixture was agitated at 1000 rpm for 10 min at 4°C. The extracts were subsequently centrifugated (16,000 xg; 4°C; 15 min). The supernatants were collected and transferred to new tubes, lyophilized, and stored at −20°C until further analysis. The samples were resuspended in 30 μl 0.1% formic acid of which 8 μl were transferred to a pooled sample for quality control (QC). 5 µl were injected using a Vanquish Horizon UPLC (Thermo Fisher Scientific; Waltham, MA) and compounds were separated on a Zorbax Eclipse Plus C18 guard (2.1 × 50 mm and 1.8 μm particle size; Agilent Technologies; Santa Clara, CA), and an analytical column (2.1 × 150 mm and 1.8 μm particle size; Agilent Technologies; Santa Clara, CA) kept at 40°C. The analytes were eluted using a flow rate of 400 μl/min and the following composition of eluent A (0.1% formic acid) and eluent B (0.1% formic acid; acetonitrile) solvents: 3% B from 0 to 1.5 min, 3% to 40% B from 1.5 to 4.5 min, 40% to 95% B from 4.5 to 7.5 min, 95% B from 7.5 to 10.1 min, and 95% to 3% B from 10.1 to 10.5 min before equilibration for 3.5 min with the initial conditions. The flow from the UPLC was coupled to a Q Exactive HF mass spectrometer (Thermo Fisher Scientific; Waltham, MA) for mass spectrometric analysis in both positive and negative ion mode.

Raw data was processed with MzMine^79^ (v 2.53). Compounds were annotated at metabolomics standards initiative^80^ level 2 using local MS/MS spectra databases of National Institute of Standards and Technology 17 (NIST17) and MassBank of North America (MoNA). The datasets were corrected for signal drift using the R package statTarget.^81^ Finally, data was normalized using QC group, log transformation (base 10), and auto scaling, all done in MetaboAnalyst^82^ (https://www.metaboanalyst.ca/).

### *In silico* predictions

Changes in SRE activities were predicted using HEXplorer^25^ (https://www2.hhu.de/rna/html/hexplorer_score.php) and HExoSplice^26^ (http://bioinfo.univ-rouen.fr/HExoSplice/inputs.php). Location and changes of SR protein binding motifs were predicted using ESEfinder 3.0^83,84^ (http://krainer01.cshl.edu/cgi-bin/tools/ESE3/esefinder.cgi?process=home) with default threshold values, and changes of putative hnRNP A1 binding motifs were predicted using DeepCLIP^27^ (https://deepclip-web.compbio.sdu.dk/) with the HNRNPA1_BRUUN pre-trained model. Splice site scores were estimated as the MaxEnt score using MaxEntScan^85^ (http://genes.mit.edu/burgelab/maxent/Xmaxentscan_scoreseq.html).

## Data availability

All relevant data supporting the findings of this study are presented in the article or in supplemental material. Raw data from RNA sequencing and metabolomics will be deposited in public repositories upon acceptance of the manuscript. Additional information and other raw data are available from the corresponding author upon reasonable request.

## Acknowledgements

We thank Professor Magdalena Ugarte for the use of patient-derived fibroblasts from the collection at CEDEM (Centro de Diagnostico de Enfermedades Moleculares). We thank Arístides López-Márquez for help with the CRISPR gene editing design and protocol. We thank Asbjørn Kofoed Seide, Anne Sofie Eybye Nielsen, Michael Silva, and Stefanie Bundgaard Flyvbjerg for technical support.

This work was supported by grants to B.S.A. from the Danish Medical Research Council (no. 9039-00281B) and the Novo Nordisk Foundation (NNF 19OC0058588), and to L.R.D.: Grant PID2019-105344RB-I00/AEI/10.13039/501100011033 from the Spanish Ministry of Science and Innovation and the European Regional Development Fund, and grant (XX National Call 2020) from Fundación Ramón Areces.

## Author contributions

B.S.A. conceptualized this study. U.S.S.P. and M.D. performed minigene experiments, L.L.H. performed SPRi, M.D. and U.S.S.P. performed RNA affinity pulldown, M.R.L. performed LC-MS/MS protein quantification, M.D. performed hnRNP A1/A2 knockdown, U.S.S.P. performed CRISPR gene editing, U.S.S.P. performed SSO transfections of CRISPR-edited HepG2 cells, U.S.S.P. performed RNA sequencing and analysis with supervision from T.K.D., J.F.H. performed metabolomics, A.M. and M.D. performed SSO transfections of patient-derived fibroblasts, A.M. performed PCCA/PCCB western blot analyses and PCC enzyme activity assays. B.S.A., L.R.D., E.R., and N.J.F. provided supervision. U.S.S.P., M.D., and B.S.A. drafted and revised the manuscript. All authors reviewed the final draft.

