## Supplemental material for "Regulating *PCCA* gene expression by modulation of pseudoexon splicing patterns to rescue enzyme activity in propionic acidemia"

**Table S1** List of 65 possible off-targets of SSO 2 (WT; Table S2). This includes the top 50 possible off-targets with 100% identity and all possible off-targets with <100% identity in 5000 maximum target sequences from NCBI Nucleotide BLAST (https://blast.ncbi.nlm.nih.gov/Blast.cgi) of the SSO target sequence, with nucleotide collection (nr/nt) as selected database for Homo sapiens (taxid:9606) and program selection optimized for somewhat similar sequences (blastn) with default settings. The number of identical nucleotides between the target sequence and possible off-targets is indicated as identity, and exon/intron location of possible off-targets is given corresponding to the relative RefSeq sequences (MANE Select). Intronic locations <500 bp from exons are indicated.

| Gene | RefSeq | Location | Identity (nt) |
|---|---|---|---|
| ***FGF14*** | NM_004115 | Intron 3 | 17/17 |
| ***NTRK2*** | NM_006180 | Exon 19 (3’UTR) | 16/16 |
| ***GRIP1*** | NM_001366722 | Upstream first exon | 16/16 |
| ***PRUNE2*** | NM_015225 | Exon 8 | 16/16 |
| ***MBD5*** | NM_001378120 | Intron 1 | 16/16 |
| ***KLRC1*** | NM_002259 | Intron 5<br>(478 bp downstream exon 5) | 16/16 |
| ***MACROD2*** | NM_001351661 | Intron 2<br>Intron 8 | 15/15<br>15/15 |
| ***HERC2*** | NM_004667 | Intron 76<br>(150 bp downstream exon 76;<br>226 bp upstream exon 77) | 15/15 |
| ***RHCE*** | NM_020485 | Intron 9 | 15/15 |
| ***RYR3*** | NM_001036 | Intron 93 | 15/15 |
| ***HDAC9*** | NM_178425 | Upstream first exon | 15/15 |
| ***SAMD5*** | NM_001030060 | [a] | 15/15 |
| ***GRM7*** | NM_000844 | Intron 8 | 15/15 |
| ***TFCP2*** | NM_005653 | Intron 10<br>(131 bp downstream exon 10) | 15/15 |
| ***PCDH11X*** | NM_032968 | Intron 9 | 15/15 |
| ***PCDH11Y*** | NM_001395587 | Intron 4<br>Intron 3 | 15/15<br>16/17 |
| ***SGCZ*** | NM_139167 | Intron 1 | 15/15 |
| ***RHD*** | NM_016124 | Intron 9 | 15/15 |
| ***DNAAF11*** | NM_012472 | Exon 4 | 14/14 |
| ***RABGAP1L*** | NM_001366446 | Exon 26<br>(3’UTR) | 14/14 |
| ***MSL1*** | NM_001365919 | Exon 9<br>(3’UTR) | 14/14 |
| ***TPK1*** | NM_022445 | Intron 7 | 14/14 |
| ***MECOM*** | NM_004991 | Intron 5 | 14/14 |
| ***RELN*** | NM_005045 | Intron 2 | 14/14 |
| ***RPE65*** | NM_000329 | Downstream last exon | 14/14 |
| ***GRIN2A*** | NM_001134407 | Intron 7<br>(3 bp downstream exon 7) | 14/14 |
| ***CTNND2*** | NM_001332 | Intron 9 | 14/14 |
| ***CYP2C9*** | NM_000771 | Intron 5 | 14/14 |
| ***CYP2C19*** | NM_000769 | Intron 5 | 14/14 |
| ***SLC25A26*** | NM_001379210 | Downstream last exon<br>Intron 4<br>(59 bp downstream exon 4) | 14/14<br>16/17 |
| ***TBL1X*** | NM_005647 | Intron 2 | 14/14 |
| ***ABLIM1*** | NM_002313 | Upstream first exon | 14/14 |
| ***GPSM2*** | NM_013296 | Intron 5<br>(273 bp downstream exon 5;<br>253 bp upstream exon 6) | 14/14 |

| | | | |
|---|---|---|---|
| ***TENM4*** | NM_001098816 | Intron 3<br>Intron 6 | 14/14<br>17/18 |
| ***ADGRV1*** | NM_032119 | Intron 1 | 14/14 |
| ***LRP1B*** | NM_018557 | Intron 3 | 14/14 |
| ***EXOC6B*** | NM_015189 | Intron 6 | 14/14 |
| ***AGMO*** | NM_001004320 | Intron 12[a] | 14/14 |
| ***CNTN5*** | NM_014361 | Intron 16 | 14/14 |
| ***UBR5*** | NM_015902 | Intron 1 | 14/14 |
| ***PCLO*** | NM_033026 | Intron 9 | 14/14 |
| ***MED13*** | NM_005121 | Intron 8 | 14/14 |
| ***ZNF804A*** | NM_194250 | Intron 1 | 14/14 |
| ***HIVEP2*** | NM_006734 | Intron 2 | 14/14 |
| ***AASDH*** | NM_181806 | Intron 1 | 14/14 |
| ***EPHX1*** | NM_001136018 | Intron 4<br>(255 bp downstream exon 4;<br>66 bp upstream exon 5) | 14/14 |
| ***SIPA1L3*** | NM_015073 | Intron 13 | 14/14 |
| ***PSPC1*** | NM_001354909 | Downstream last exon[a] | 14/14 |
| ***LYST*** | NM_000081 | Intron 31[a] | 14/14 |
| ***MIPOL1*** | NM_001388067 | Intron 12[a] | 14/14 |
| ***POU6F2*** | NM_001370959 | Intron 4 | 17/18 |
| ***STIM1*** | NM_001382567 | Intron 1 | 17/18 |
| ***KAZN*** | NM_201628 | Upstream first exon | 16/17 |
| ***ANO6*** | NM_001025356 | Exon 20<br>(3'UTR) | 16/17 |
| ***TTC30B*** | NM_152517 | Exon 1/1 | 16/17 |
| ***TTF2*** | NM_003594 | Exon 23<br>(3'UTR) | 16/17 |
| ***TPX2*** | NM_012112 | Exon 12 | 16/17 |
| ***RBFOX1*** | NM_018723 | Intron 1 | 16/17 |
| ***TCF12*** | NM_207037 | Intron 9<br>(368 bp downstream exon 9) | 16/17 |
| ***BNC2*** | NM_017637 | Intron 5 | 16/17 |
| ***NRP1*** | NM_003873 | Intron 10<br>(274 bp downstream exon 10) | 16/17 |
| ***KLRC2*** | NM_002260 | Intron 10<br>(475 bp downstream exon 4) | 16/17 |
| ***SLC17A8*** | NM_139319 | Intron 9<br>(2 bp downstream exon 9) | 16/17 |
| ***CD3D*** | NM_000732 | Downstream last exon | 16/17 |
| ***TEAD1*** | NM_021961 | Intron 10 | 19/21 |

[a] Predicted in another mRNA transcript variant.

**Table S2** List of splice-switching antisense oligonucleotides (SSO). All SSOs used are RNA oligonucleotides with 2’-O-methyl modifications and phosphorothioate backbones (LGC Biosearch Technologies; Risskov, Denmark).

| **ID** | **Sequence (5’-3’)** |
|---|---|
| **SSO 1** (WT)[a] | UUCUAUGAAAUUCACAGUGUUGACA |
| **SSO 2** (WT)[a] | AUGUUCUAUGAAAUUCACAGUGUUG |
| **SSO 3** (WT)[a] | ACUGAUGUUCUAUGAAAUUCACAGU |
| **SSO 4** (WT)[a] | UAAACUGAUGUUCUAUGAAAUUCAC |
| **SSO 5** (WT)[a] | ACAUAAACUGAUGUUCUAUGAAAUU |
| **SSO 6** (WT)[a] | AGGACAUAAACUGAUGUUCUAUGAA |
| **SSO 7** (WT)[a] | UUCAGGACAUAAACUGAUGUUCUAU |
| **Ctrl SSO** (25-mer)[a] | GCUCAAUAUGCUACUGCCAUGCUUG |
| **SSO 2** (MUT)[b] | AUGUUCCAUGAAAUUCACAGUGUUG |
| **Ctrl SSO** (22-mer)[b] | CAAUAUGCUACUGCCAUGCUUG |

[a] Used in transfection of HepG2 cells
[b] Used in transfection of fibroblasts.

**Table S3** List of biotinylated RNA oligonucleotides (LGC Biosearch Technologies; Risskov, Denmark).

| ID | Sequence (5'-3') |
|---|---|
| **WT 1** | AAUUUCAUAGAACAUCAGU-biotin |
| **MUT 1** | AAUUUCAUGGAACAUCAGU-biotin |
| **WT 2** | GUGAAUUUCAUAGAACAUCAGU-biotin |
| **MUT 2** | GUGAAUUUCAUGGAACAUCAGU-biotin |
| **WT** | UUUUCAGAAAATAGAAAGCCAAGUA-biotin |
| **M3A1** | UUUUCCGAAAATAGAAAGCCAAGUA-biotin |
| **M4A1** | UUUUCAGAAAAUCGAAAGCCAAGUA-biotin |
| **M3A1+M4A1** | UUUUCCGAAAAUCGAAAGCCAAGUA-biotin |

**A**

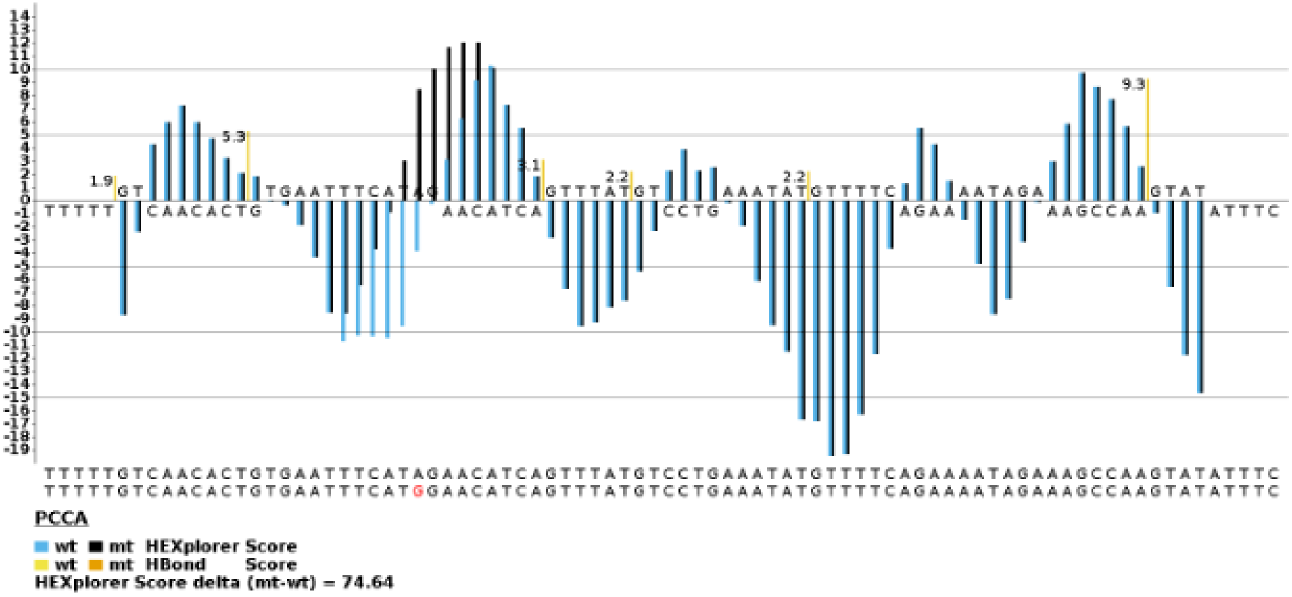


**B**

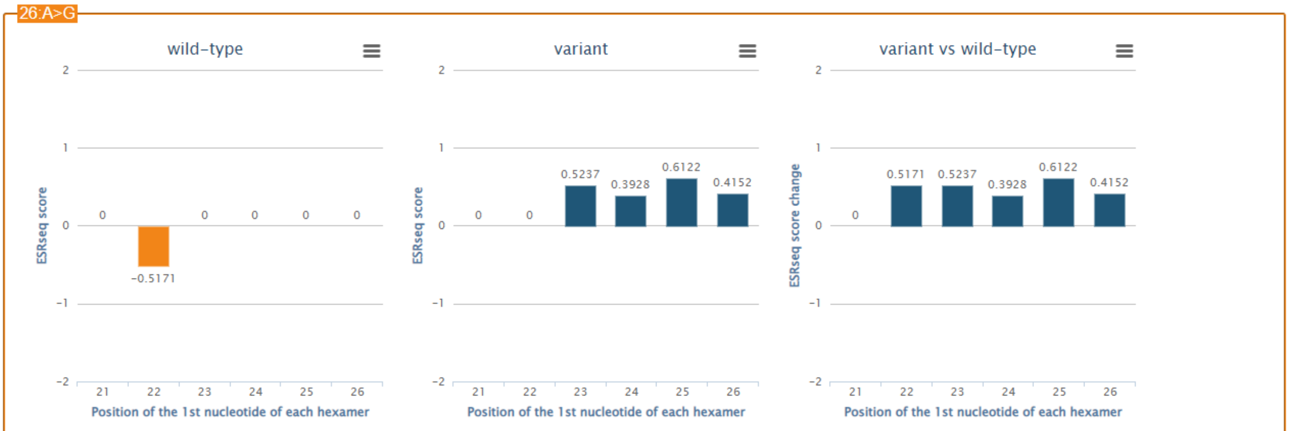


| Positions | Hexamer seq wild-type | ESRseq score wild-type | Hexamer assignment wild-type | Hexamer seq variant | ESRseq score variant | Hexamer assignment variant | ESRseq score change (variant vs wild-type) |
|---|---|---|---|---|---|---|---|
| 21-26 | TTCAT**A** | 0 | N | TTCAT**G** | 0 | N | 0 |
| 22-27 | TCAT**A**G | -0.5171 | S | TCAT**G**G | 0 | N | +0.5171 |
| 23-28 | CAT**A**GA | 0 | N | CAT**G**GA | +0.5237 | E | +0.5237 |
| 24-29 | AT**A**GAA | 0 | N | AT**G**GAA | +0.3928 | E | +0.3928 |
| 25-30 | T**A**GAAC | 0 | N | T**G**GAAC | +0.6122 | E | +0.6122 |
| 26-31 | **A**GAACA | 0 | N | **G**GAACA | +0.4152 | E | +0.4152 |
| Total | TTCAT**A**GAACA | -0.5171 | - | TTCAT**G**GAACA | +1.9439 | - | +2.4610 |

Download table

**Figure S1** Analysis of the splicing regulatory change in the *PCCA* pseudoexon caused by the c.1285-1416A>G sequence variation. **(A)** Analysis by HEXplorer[1] (https://www2.hhu.de/rna/html/hexplorer_score.php), with the A>G variation marked in red. **(B)** Analysis by HExoSplice[2] (http://bioinfo.univ-rouen.fr/HExoSplice/inputs.php), with the A>G variation located at position 26 in the pseudoexon sequence. Both indicates that the *PCCA* pseudoexon activating variation changes the adjacent splicing regulatory properties from silencing to enhancing, by changing the relative scores from negative to positive (HEXplorer $HZ_{EI}$ scores for each nucleotide between wild type marked in blue and variant marked in black, and HExoSplice ESRseq scores for overlapping hexamer sequences).

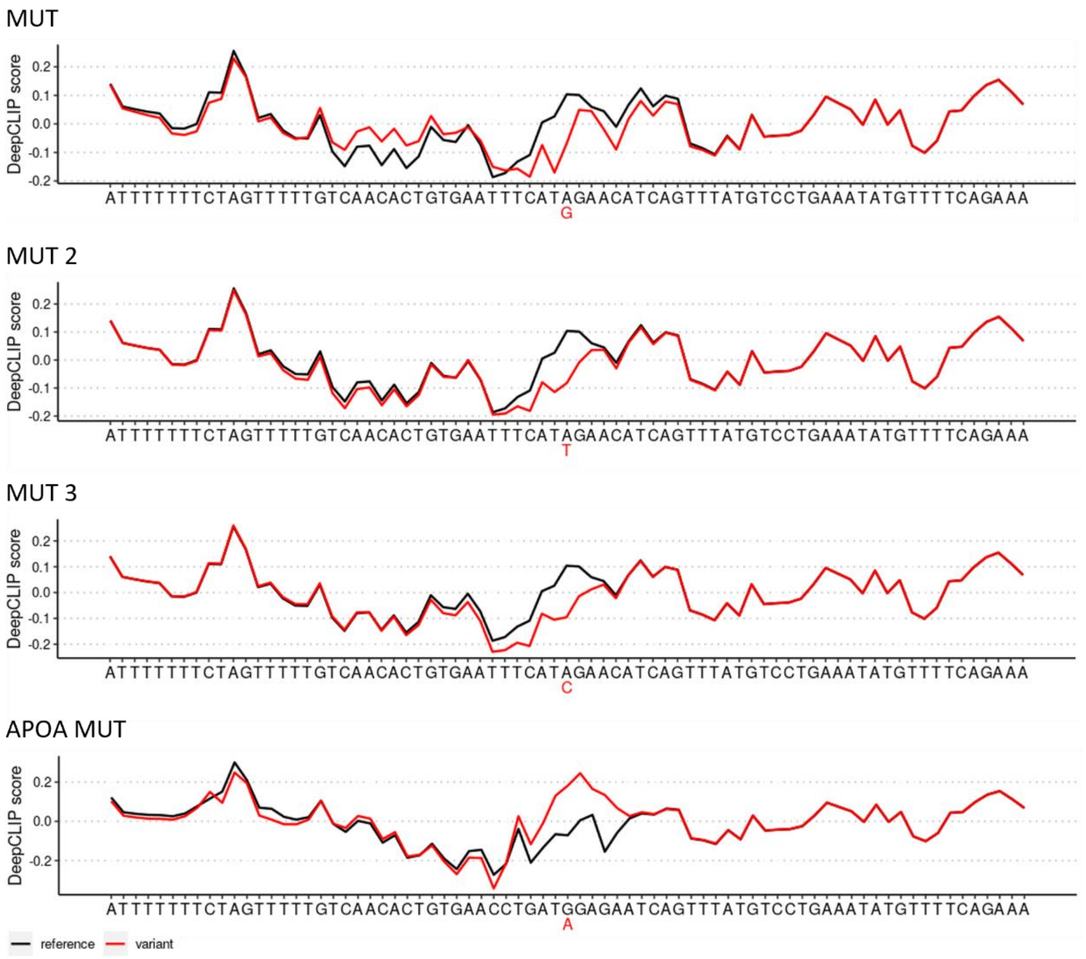


**Figure S2** Changes in predicted binding profiles for hnRNP A1. This is predicted using DeepCLIP[3] (https://deepclip-web.compbio.sdu.dk/) with the HNRNPA1_BRUUN pretrained model and 75 bp sequences (-37/+37 around the variation). The binding profiles predict the MUT, MUT 2, and MUT 3 variants from our splicing reporter minigene assay (Figure 1A) against the relative WT sequence. The APOA MUT variant is predicted against the APOA WT sequence. Variant binding profiles are indicated in red.

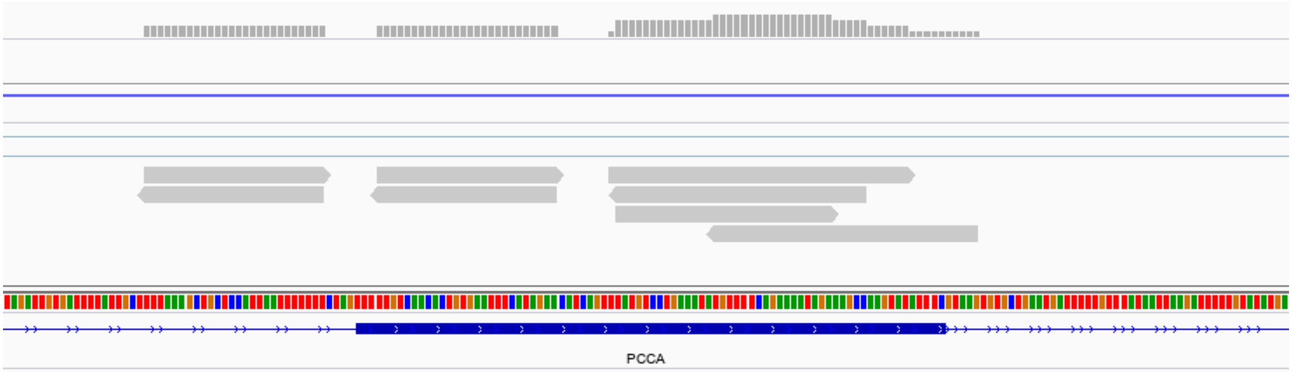


**Figure S3** Enhanced UV crosslinking and immunoprecipitation (eCLIP) reads from an hnRNP A1 target experiment in HepG2 cells. Data is downloaded from the ENCODE portal[4,5] (https://www.encodeproject.org/; replicate 1 from experiment ENCSR769UEW; bam file: ENCFF330OFU) and visualized with the Integrative Genomics Viewer (IGV)[6] for -50/+50 bp of the *PCCA* pseudoexon sequence (13:100305701-100305884; GRCh38). The *PCCA* pseudoexon is marked by the blue box.

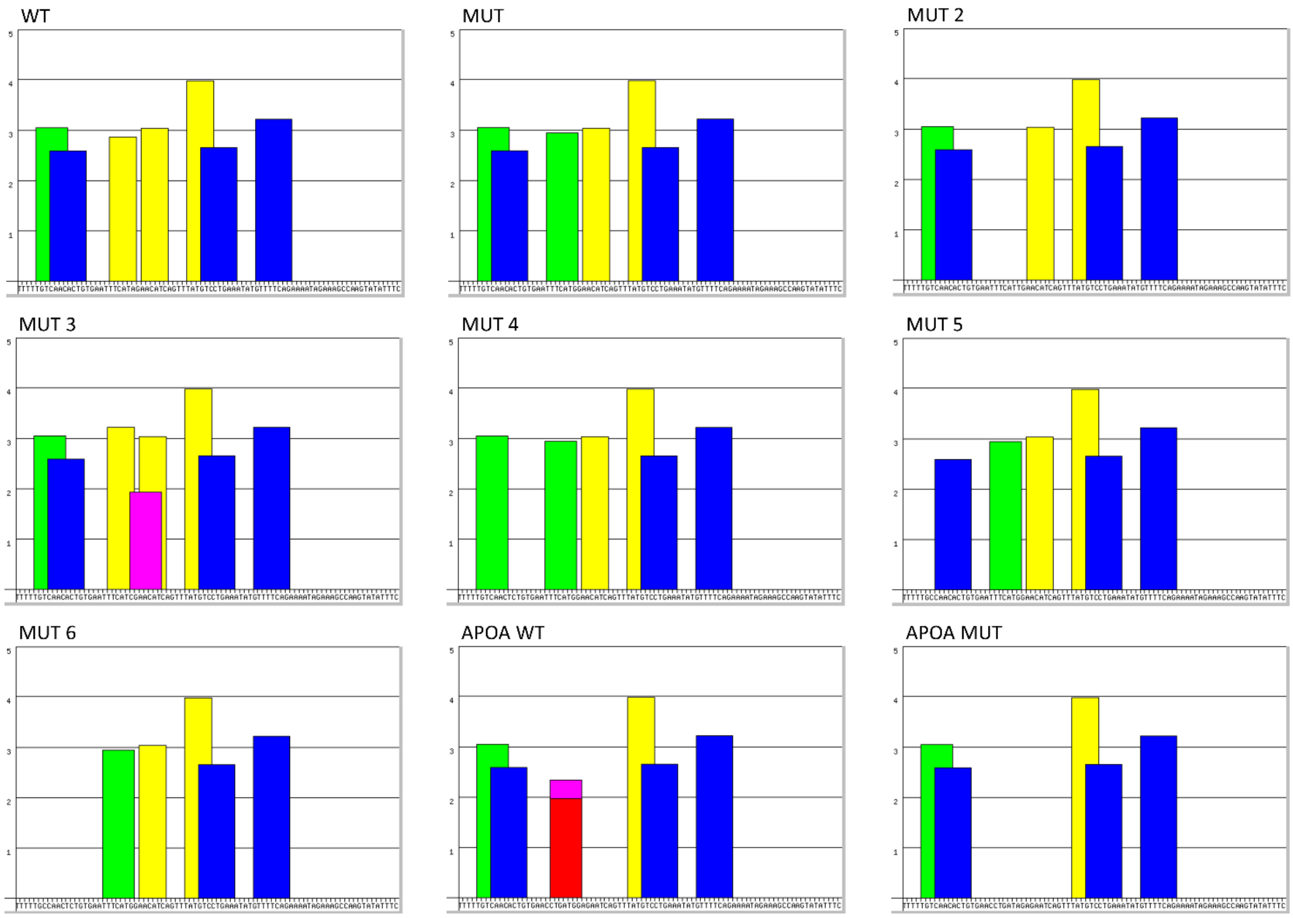


**Figure S4** Exonic splicing enhancers (ESE) predicted across variants of the *PCCA* pseudoexon sequence. The ESEs are predicted as binding motifs for SRSF1 (red/pink), SRSF2 (blue), SRSF5 (green), and SRSF6 (yellow) using ESEfinder 3.0[7,8] (http://krainer01.cshl.edu/cgi-bin/tools/ESE3/esefinder.cgi?process=home) for the WT, MUT, MUT 1-6, APOA WT, and APO MUT variants from our splicing reporter minigene assay (Figure 1A). The MUT variant disrupts a binding motif for SRSF6 and creates a binding motif for SRSF5, the MUT 2 variant disrupts the same binding motif for SRSF6, and the MUT 3 variant creates a binding motif for SRSF1 relative to WT. The MUT 4 variant disrupts a binding motif for SRSF2, the MUT 5 variant disrupts a binding motif for SRSF5, and the MUT 6 variant disrupts both the SRSF2 and SRSF5 binding motifs relative to WT. The APOA WT variant contains a predicted binding motif for SRSF1, which is disrupted by the APOA MUT variant.

A

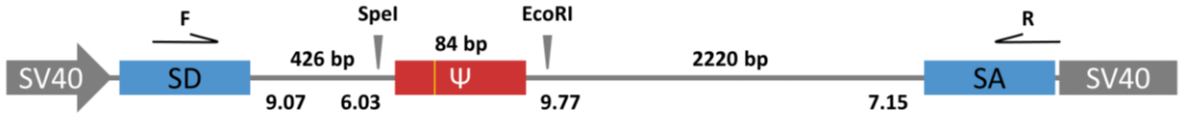


B

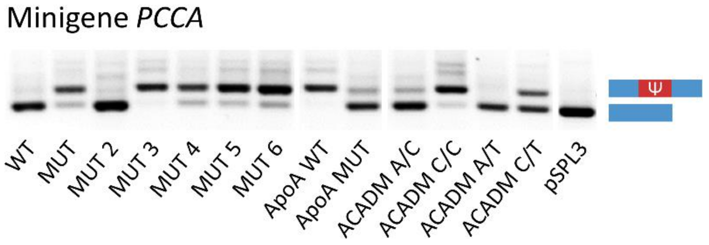


**Figure S5** Reproducing *PCCA* pseudoexon splicing patterns from minigene variants (Figure 1B) in a different splicing reporter. **(A)** Schematic representation of another *PCCA* pseudoexon splicing reporter minigene system.[9] The minigene construct contains the pseudoexon and flanking intron regions (-84/+131 bp) with an AAGGGC insert at the 3' end between SpeI and EcoRI restriction sites in the pSPL3 plasmid vector. Location of the *PCCA* c.1285-1416A>G variation is marked in yellow. Maximum entropy (MaxEnt) scores are given for each splice site in the minigene construct. F and R indicates the location of forward and reverse primers for RT-PCR analysis of the minigene pseudoexon splicing patterns. **(B)** Representative gel picture of *PCCA* pseudoexon splicing patterns from minigene variants (pSPL3 construct) transfected into HepG2 cells. MUT, MUT 2-6, APOA WT/MUT, and ACADM A/C, C/C, A/T, and C/T variants of the minigene pseudoexon sequence in the pSPL3 construct are identical to the variants of the minigene pseudoexon sequence in the pcDNA3.1 construct (Figure 1A). Splicing patterns are investigated by amplification of the region from the pSPL3 splice donor (SD) to the pSPL3 splice acceptor (SA) (SD6: 5'-TCTGAGTCACCTGGACAACC-3' and SA2: 5'-ATCTCAGTGGTATTTGTGAGC-3'). Expected fragment lengths; 255 bp (pSPL3 SD-SA)/339 bp (pSPL3 SD-SA with pseudoexon inclusion), are represented by the lowest two bands. SV40; simian virus 40, Ψ; pseudoexon.

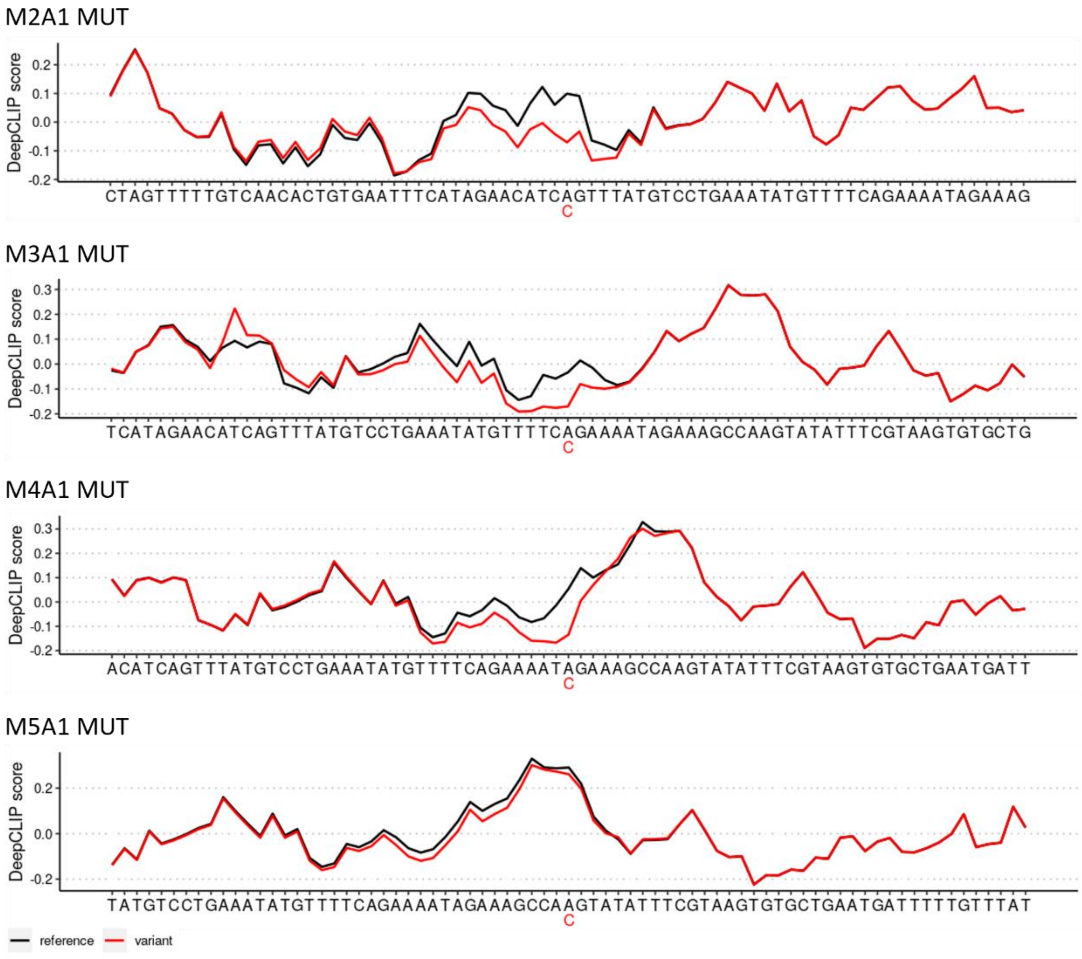


**Figure S6** Changes in predicted binding profiles for hnRNP A1. This is predicted using DeepCLIP[3] (https://deepclip-web.compbio.sdu.dk/) with the HNRNPA1_BRUUN pretrained model and 75 bp sequences (-37/+37 around the variation). The binding profiles predict the M2A1, M3A1, M4A1, and M5A1 MUT variants from our splicing reporter minigene assay (Figure 1F) against the relative WT sequence. Variant binding profiles are indicated in red.

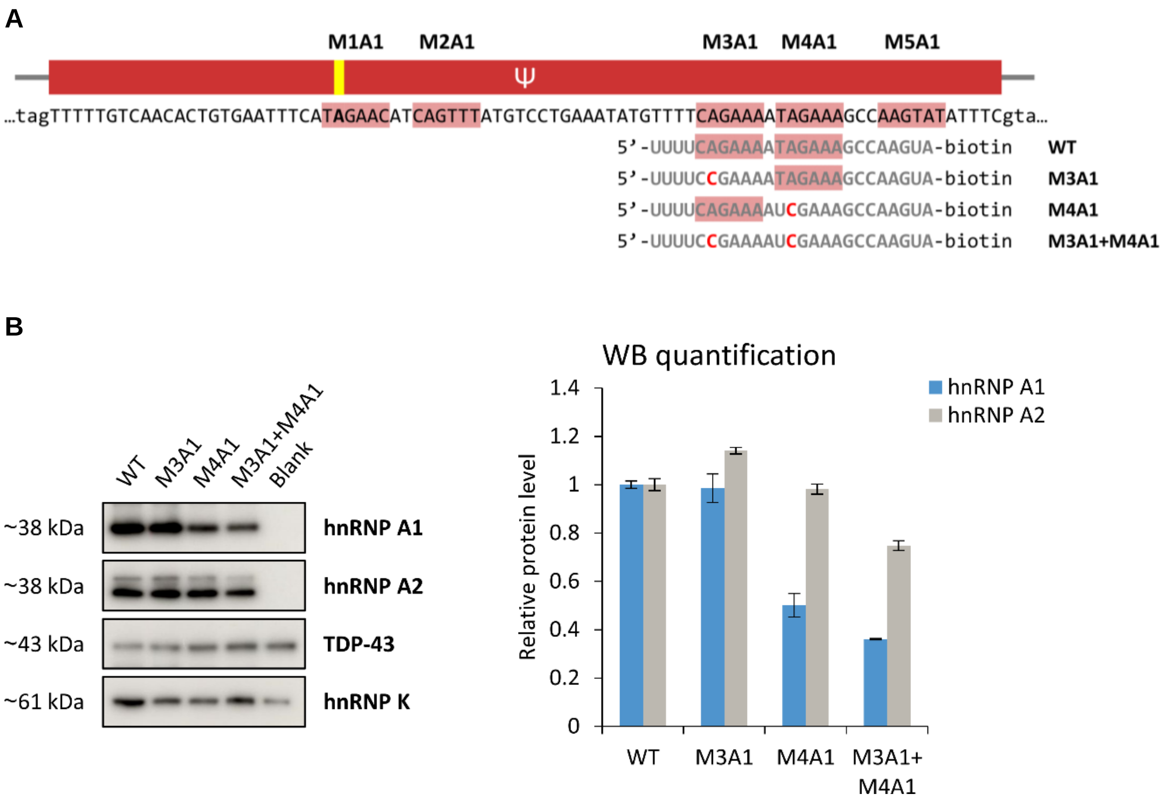


**Figure S7** Investigating binding of hnRNP A1 to another splicing regulatory region downstream of the *PCCA* c.1285-1416A>G variation. **(A)** Schematic representation and sequences of four biotinylated RNA oligonucleotides covering the region with putative hnRNP A1-binding ESSs (indicated by red highlights) used for RNA affinity pulldown with subsequent analysis by western blotting. The M3A1 and M4A1 elements are disrupted by A>C mutations of the second nucleotide in the core UAG/CAG motifs**. (B)** Representative western blots of RNA affinity pulldown samples investigating binding of hnRNP A1 (and hnRNP A2) to the wild type (WT) and mutant M3A1 and M4A1 elements of the biotinylated RNA oligonucleotides. Binding of TDP-43 and hnRNP K was used as controls illustrating different binding patterns (not much stronger than background), suggesting that the changes in binding of hnRNP A1/A2 are not due to unequal loading. Blank beads were added as a measure of background binding. Quantification by densitometry indicated as the change in protein levels relative to WT. Error bars indicate range, n=2 from individual experiments. Ψ; pseudoexon.

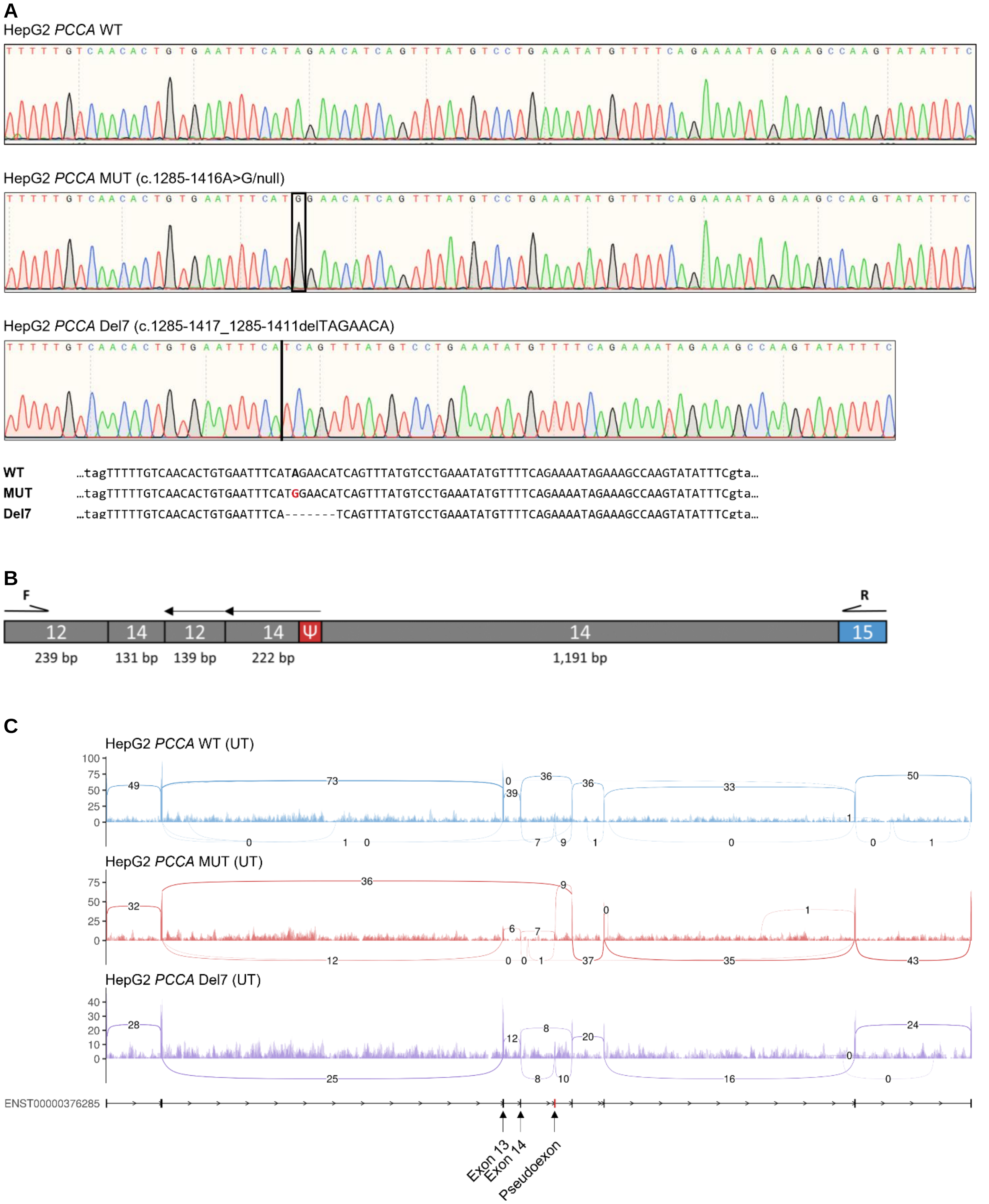


**Figure S8** Verification of the CRISPR gene editing in HepG2 cells. **(A)** Sequencing chromatograms covering the *PCCA* pseudoexon from Sanger sequencing of PCR products amplified from genomic DNA extracted from the HepG2 *PCCA* WT, MUT, and Del7 cell lines (PCCA.int14.F: 5'-TGACATAGTGGTCAAATTAGCTCT-3' and PCCA.int.14.R: 5'-CTTTGTAAGGTTGTAAAAGGAACAC-3'). **(B)** Schematic representation of region amplified between *PCCA* intron 12 and exon 15 (PCCA.int12.F: 5'-GGCCACATTGAAGATTGCCG-3' and PCCA.ex15.R) from genomic DNA extracted from the HepG2 *PCCA* MUT cell line, which revealed a 4.5 kb deletion of part of intron 12, exon 13, intron 13, exon 14, and part of intron 14, and insertion of reversed regions from intron 12 and intron 14 on one allele identified by Sanger sequencing. **(C)** Sashimi plots of the region from *PCCA* (NM_000282; Ensembl transcript ID: ENST00000376285) exon 11 to exon 18 with aggregated read coverage (y-axis) of RNA sequencing data from untreated (UT) HepG2 *PCCA* WT, MUT, and Del7 cell lines showing a lower read coverage and skipping of exon 13-14 in HepG2 *PCCA* MUT, indicating deletion of these exons on one allele, generated using ggsashimi (https://github.com/guigolab/ggsas). Ψ; pseudoexon.

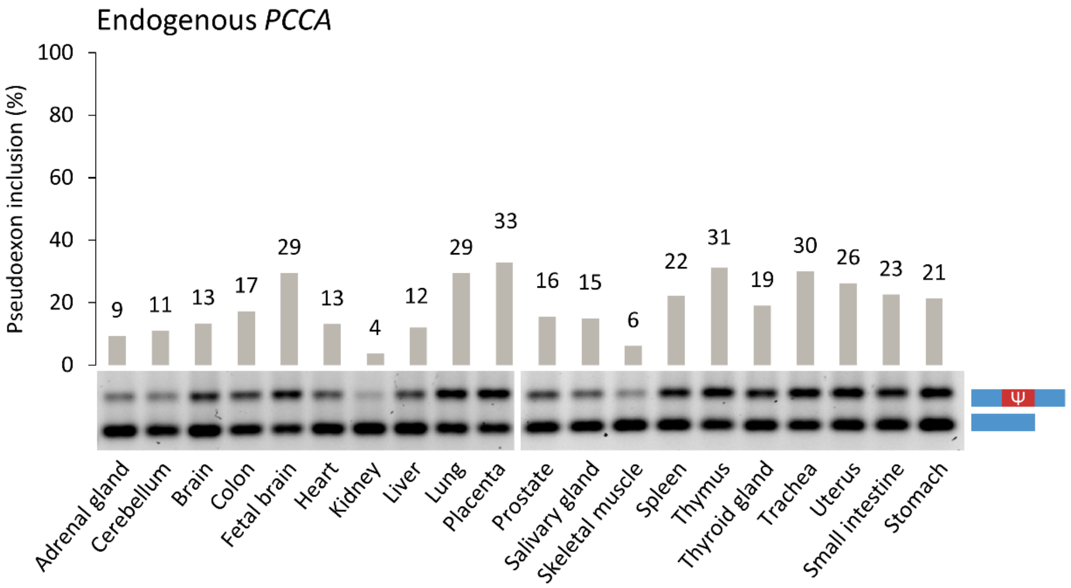


**Figure S9** Endogenous *PCCA* pseudoexon splicing patterns across a human tissue panel; Human Total RNA Master Panel II (Clontech; Mountain View, CA). Ψ; pseudoexon.

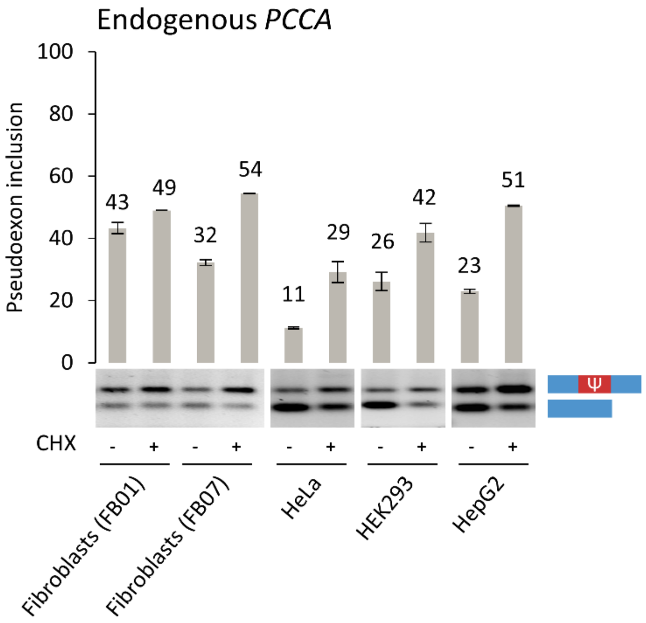


**Figure S10** Endogenous *PCCA* *p*seudoexon splicing patterns in two control fibroblast cell lines, HeLa, HEK293, and HepG2 cells, with and without cycloheximide (CHX) treatment. Error bars indicate range, n=2 culture wells. Ψ; pseudoexon.

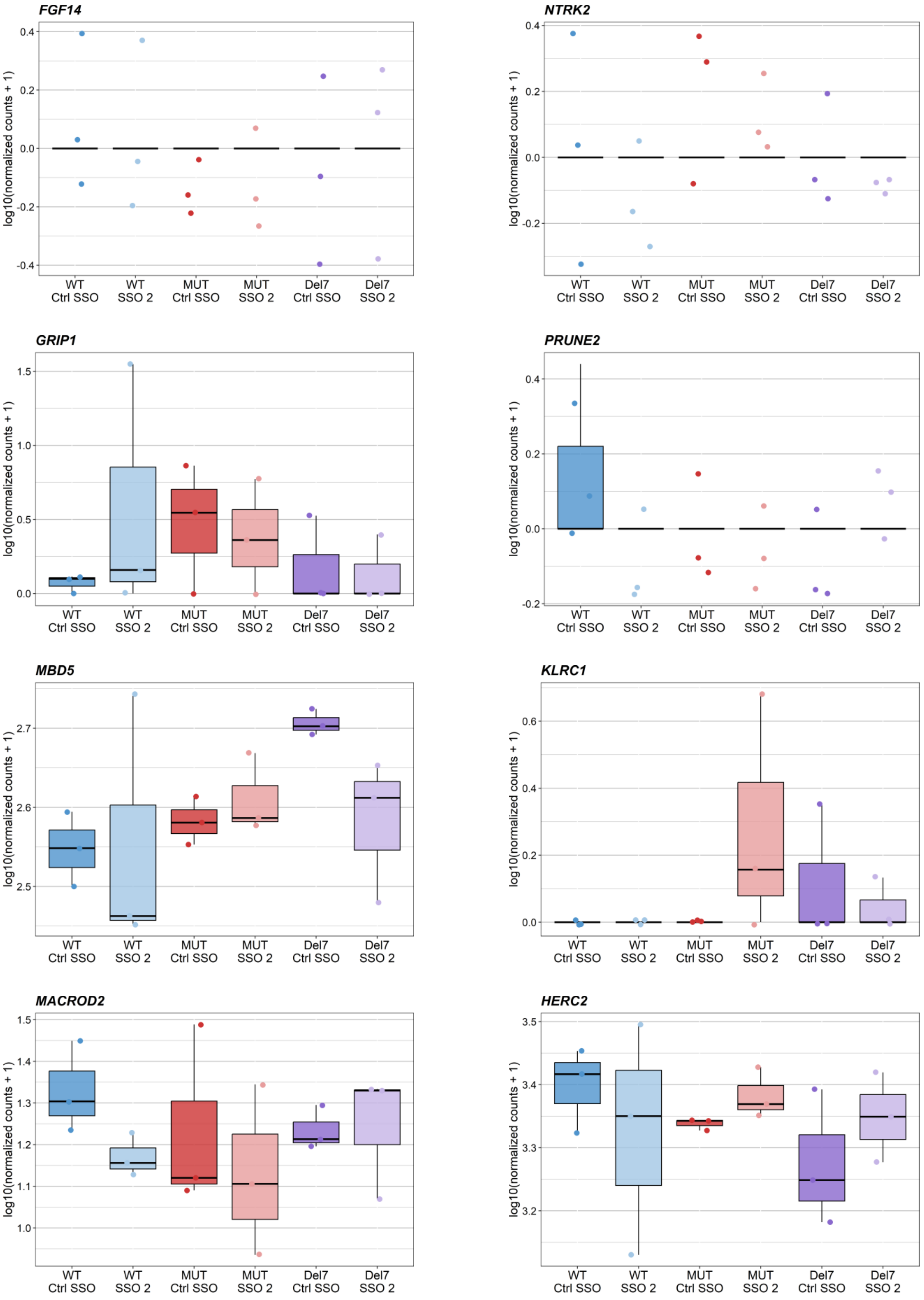
FGF14
NTRK2
GRIP1
PRUNE2
MBD5
KLRC1
MACROD2
HERC2
log10(normalized counts + 1)
WT Ctrl SSO
WT SSO 2
MUT Ctrl SSO
MUT SSO 2
Del7 Ctrl SSO
Del7 SSO 2

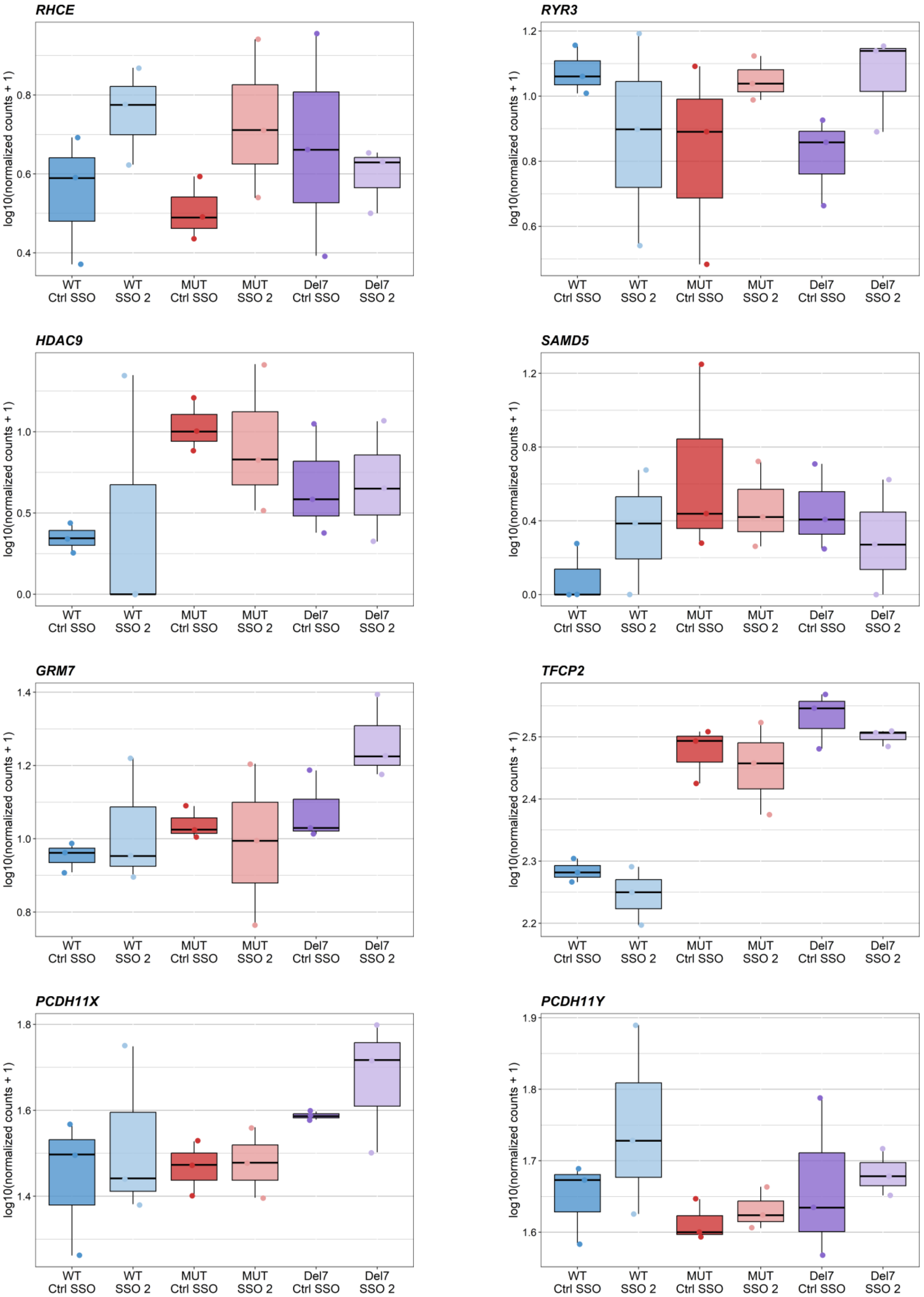

RHCE
RYR3
HDAC9
SAMD5
GRM7
TFCP2
PCDH11X
PCDH11Y
log10(normalized counts + 1)
WT Ctrl SSO
WT SSO 2
MUT Ctrl SSO
MUT SSO 2
Del7 Ctrl SSO
Del7 SSO 2

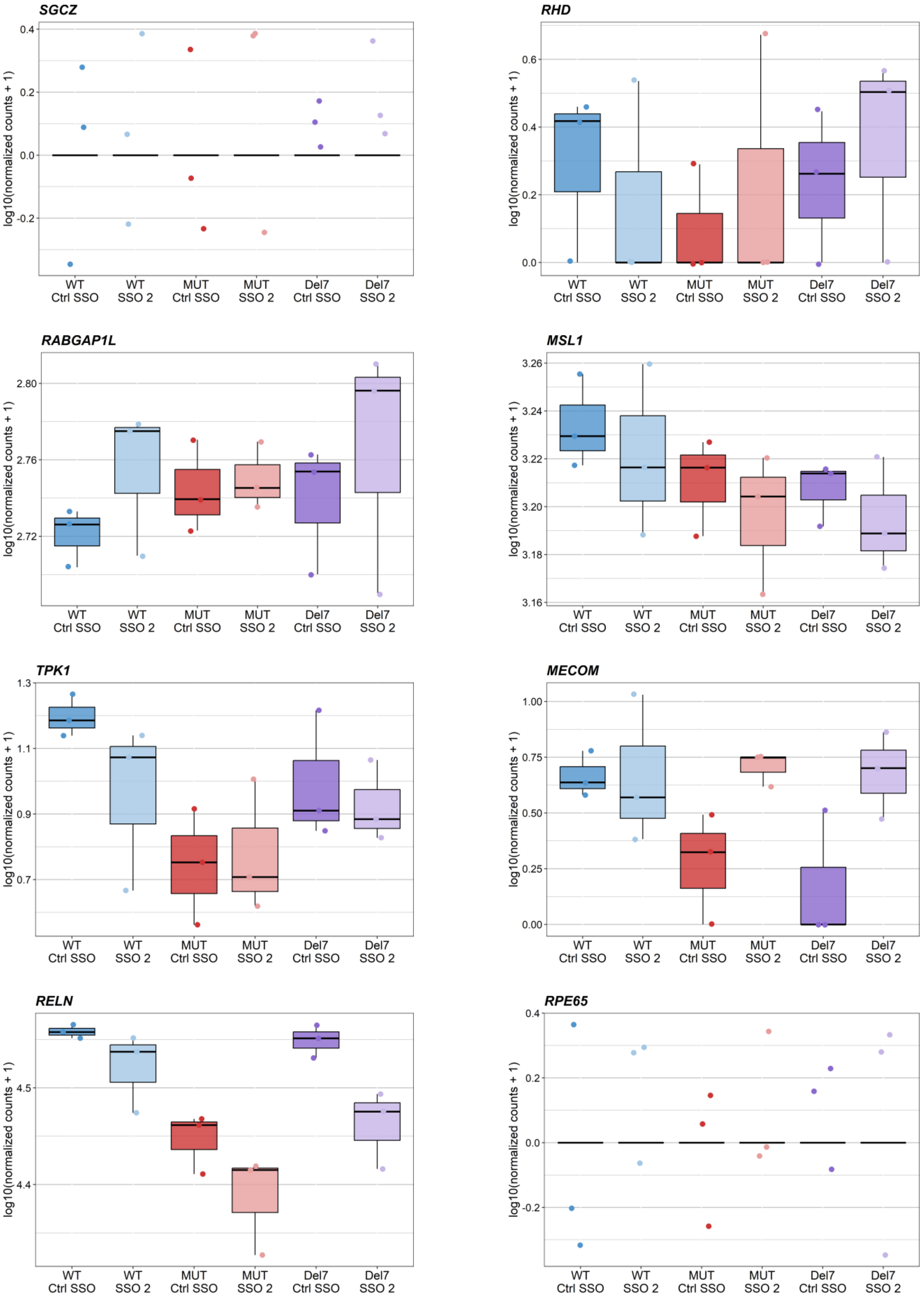
SGCZ
RHD
RABGAP1L
MSL1
TPK1
MECOM
RELN
RPE65
log10(normalized counts + 1)
WT Ctrl SSO
WT SSO 2
MUT Ctrl SSO
MUT SSO 2
Del7 Ctrl SSO
Del7 SSO 2

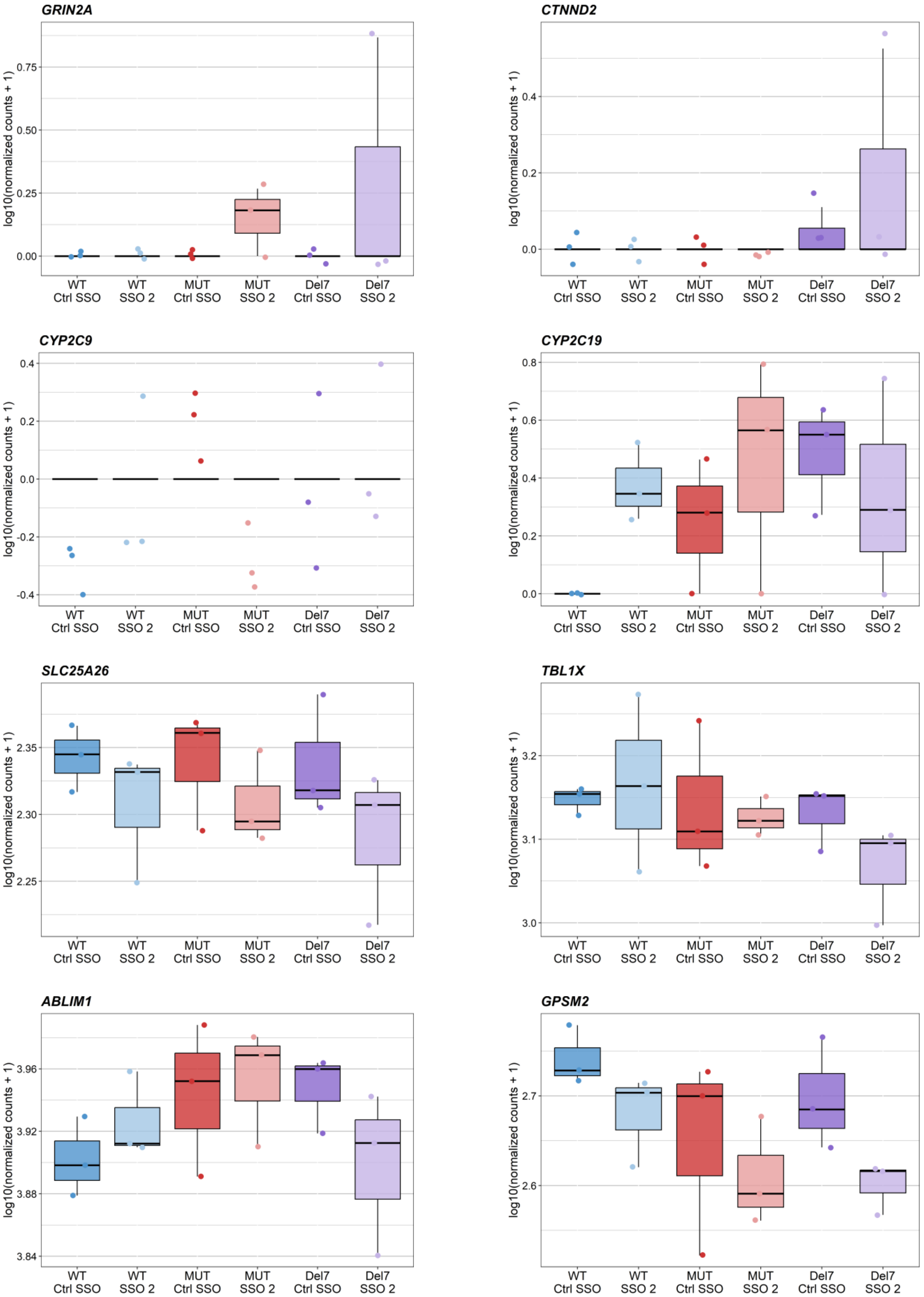
GRIN2A
CTNND2
CYP2C9
CYP2C19
SLC25A26
TBL1X
ABLIM1
GPSM2
log10(normalized counts + 1)
WT Ctrl SSO
WT SSO 2
MUT Ctrl SSO
MUT SSO 2
Del7 Ctrl SSO
Del7 SSO 2

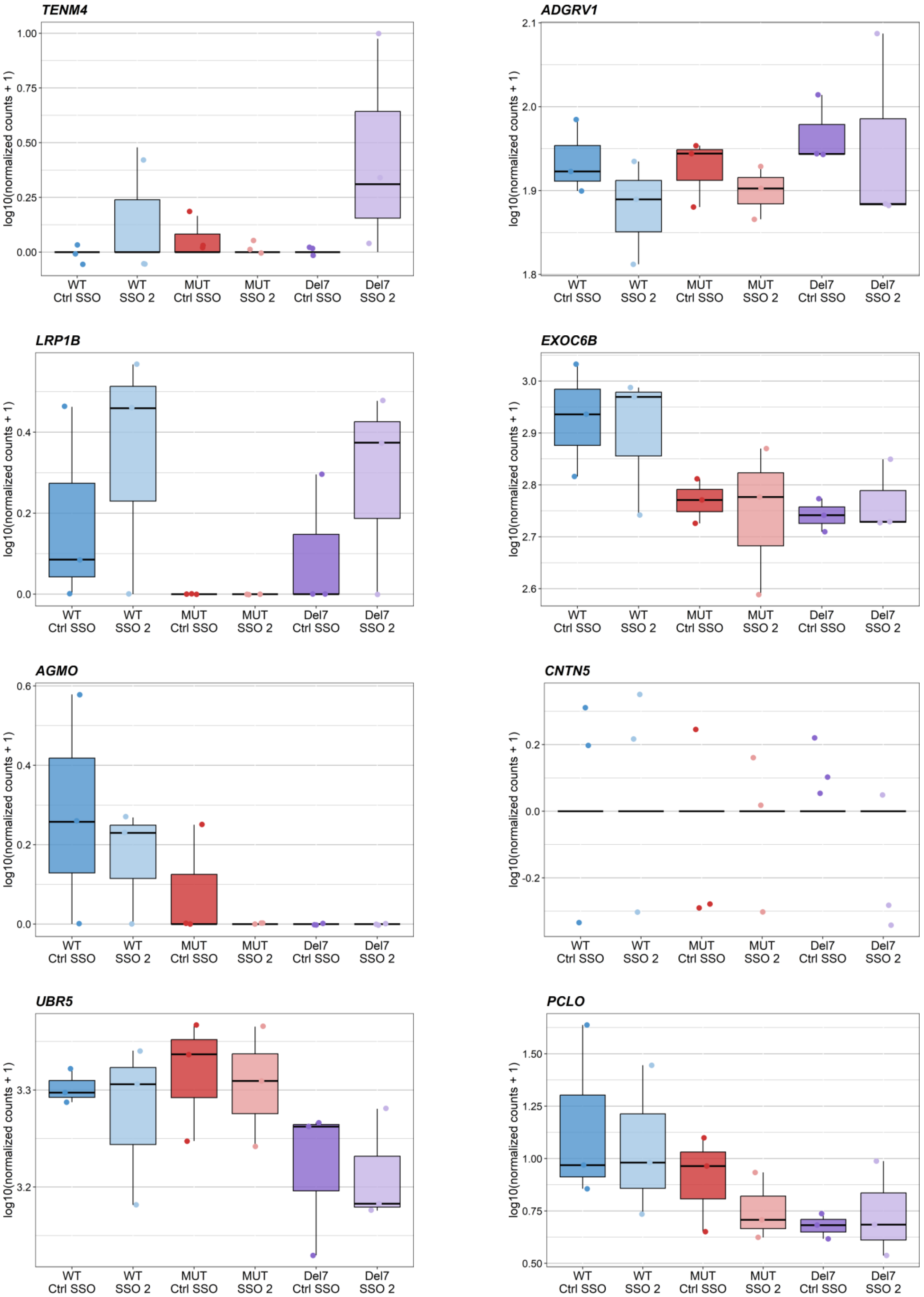
TENM4
ADGRV1
LRP1B
EXOC6B
AGMO
CNTN5
UBR5
PCLO
log10(normalized counts + 1)
WT Ctrl SSO
WT SSO 2
MUT Ctrl SSO
MUT SSO 2
Del7 Ctrl SSO
Del7 SSO 2

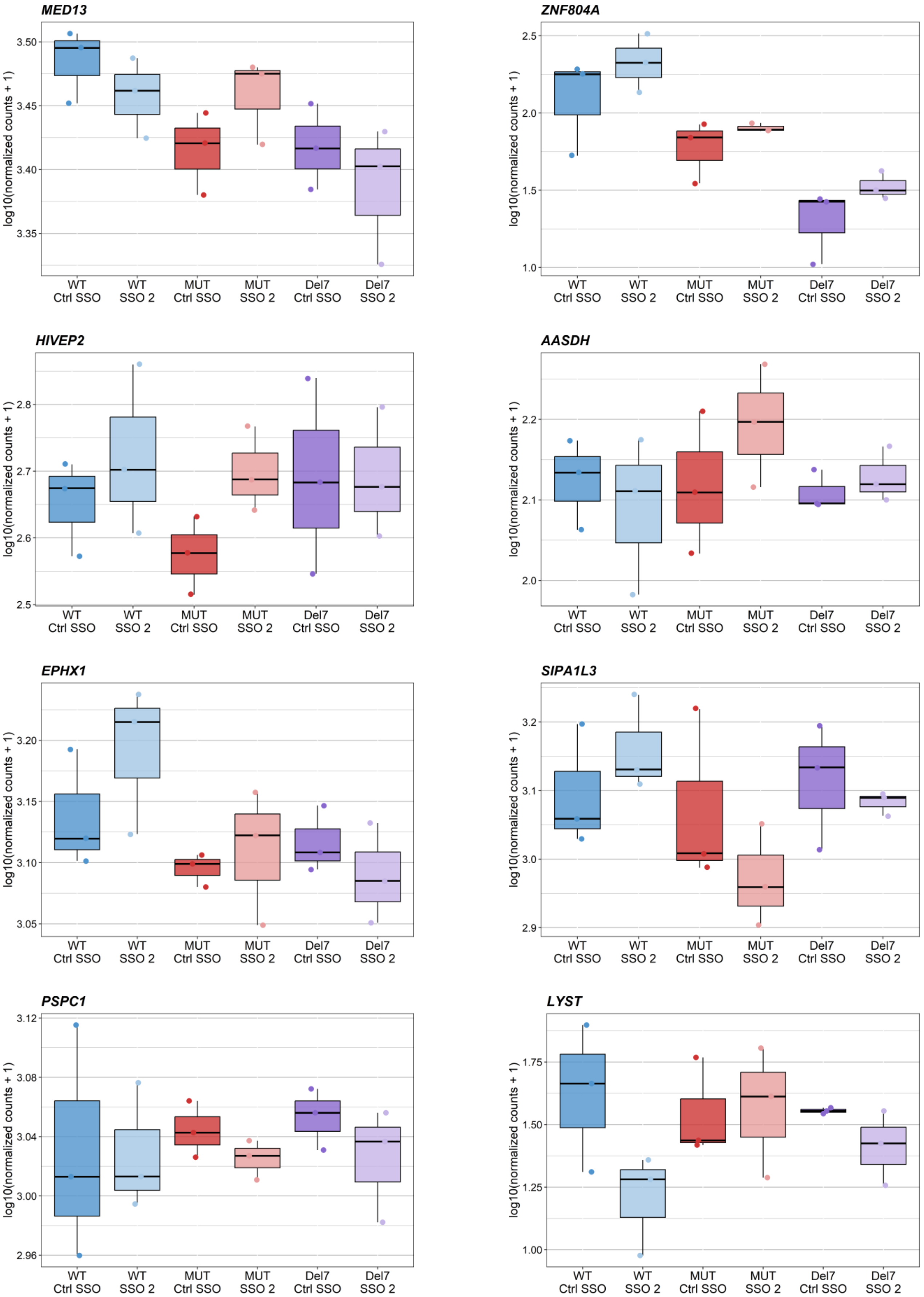
MED13
ZNF804A
HIVEP2
AASDH
EPHX1
SIPA1L3
PSPC1
LYST
log10(normalized counts + 1)
3.50
3.45
3.40
3.35
2.5
2.0
1.5
1.0
2.8
2.7
2.6
2.5
2.2
2.1
2.0
3.20
3.15
3.10
3.05
3.2
3.1
3.0
2.9
3.12
3.08
3.04
3.00
2.96
1.75
1.50
1.25
1.00
WT Ctrl SSO
WT SSO 2
MUT Ctrl SSO
MUT SSO 2
Del7 Ctrl SSO
Del7 SSO 2

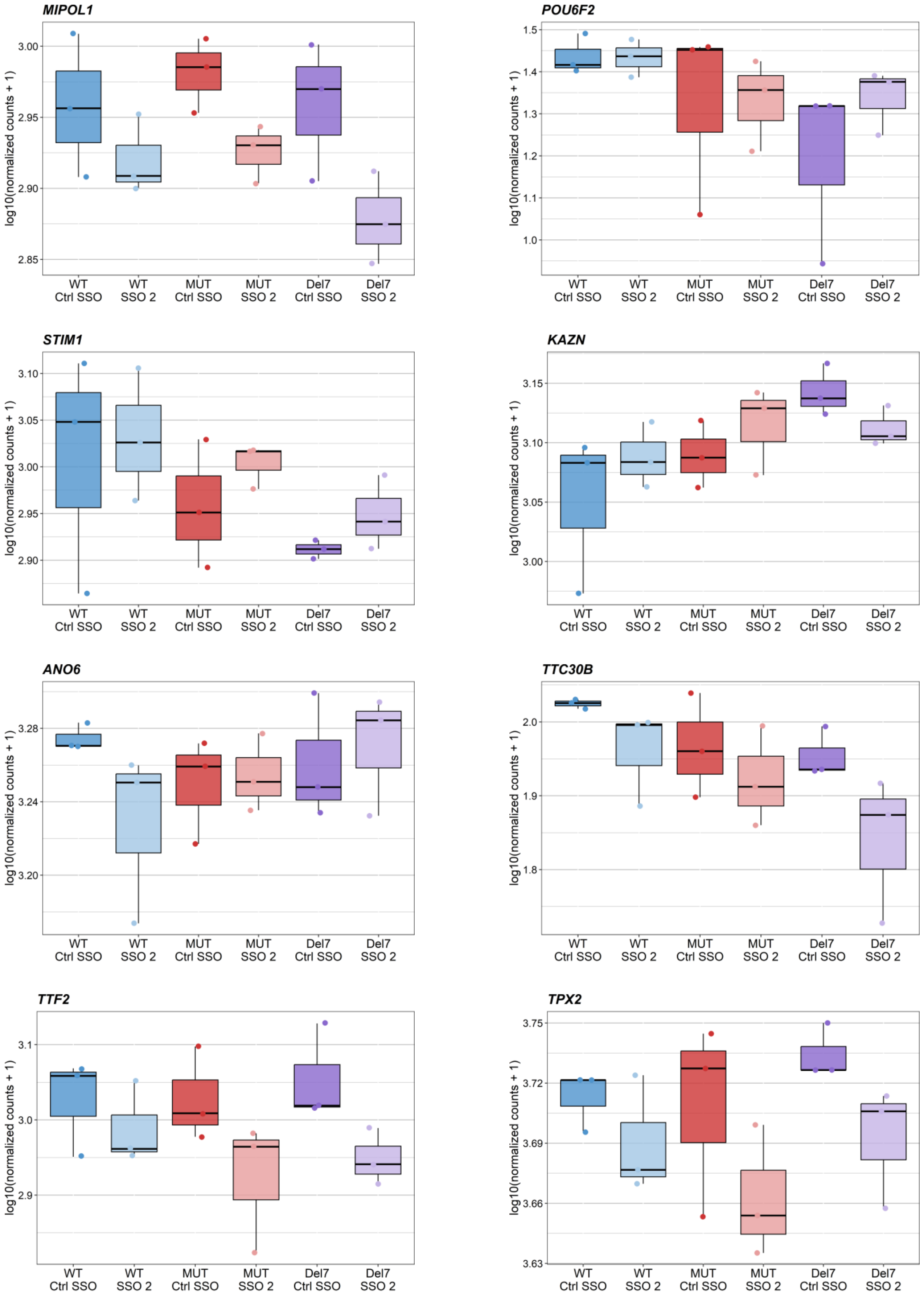
MIPOL1
log10(normalized counts + 1)
3.00
2.95
2.90
2.85
WT Ctrl SSO
WT SSO 2
MUT Ctrl SSO
MUT SSO 2
Del7 Ctrl SSO
Del7 SSO 2
POU6F2
log10(normalized counts + 1)
1.5
1.4
1.3
1.2
1.1
1.0
WT Ctrl SSO
WT SSO 2
MUT Ctrl SSO
MUT SSO 2
Del7 Ctrl SSO
Del7 SSO 2
STIM1
log10(normalized counts + 1)
3.10
3.05
3.00
2.95
2.90
WT Ctrl SSO
WT SSO 2
MUT Ctrl SSO
MUT SSO 2
Del7 Ctrl SSO
Del7 SSO 2
KAZN
log10(normalized counts + 1)
3.15
3.10
3.05
3.00
WT Ctrl SSO
WT SSO 2
MUT Ctrl SSO
MUT SSO 2
Del7 Ctrl SSO
Del7 SSO 2
ANO6
log10(normalized counts + 1)
3.28
3.24
3.20
WT Ctrl SSO
WT SSO 2
MUT Ctrl SSO
MUT SSO 2
Del7 Ctrl SSO
Del7 SSO 2
TTC30B
log10(normalized counts + 1)
2.0
1.9
1.8
WT Ctrl SSO
WT SSO 2
MUT Ctrl SSO
MUT SSO 2
Del7 Ctrl SSO
Del7 SSO 2
TTF2
log10(normalized counts + 1)
3.1
3.0
2.9
WT Ctrl SSO
WT SSO 2
MUT Ctrl SSO
MUT SSO 2
Del7 Ctrl SSO
Del7 SSO 2
TPX2
log10(normalized counts + 1)
3.75
3.72
3.69
3.66
3.63
WT Ctrl SSO
WT SSO 2
MUT Ctrl SSO
MUT SSO 2
Del7 Ctrl SSO
Del7 SSO 2

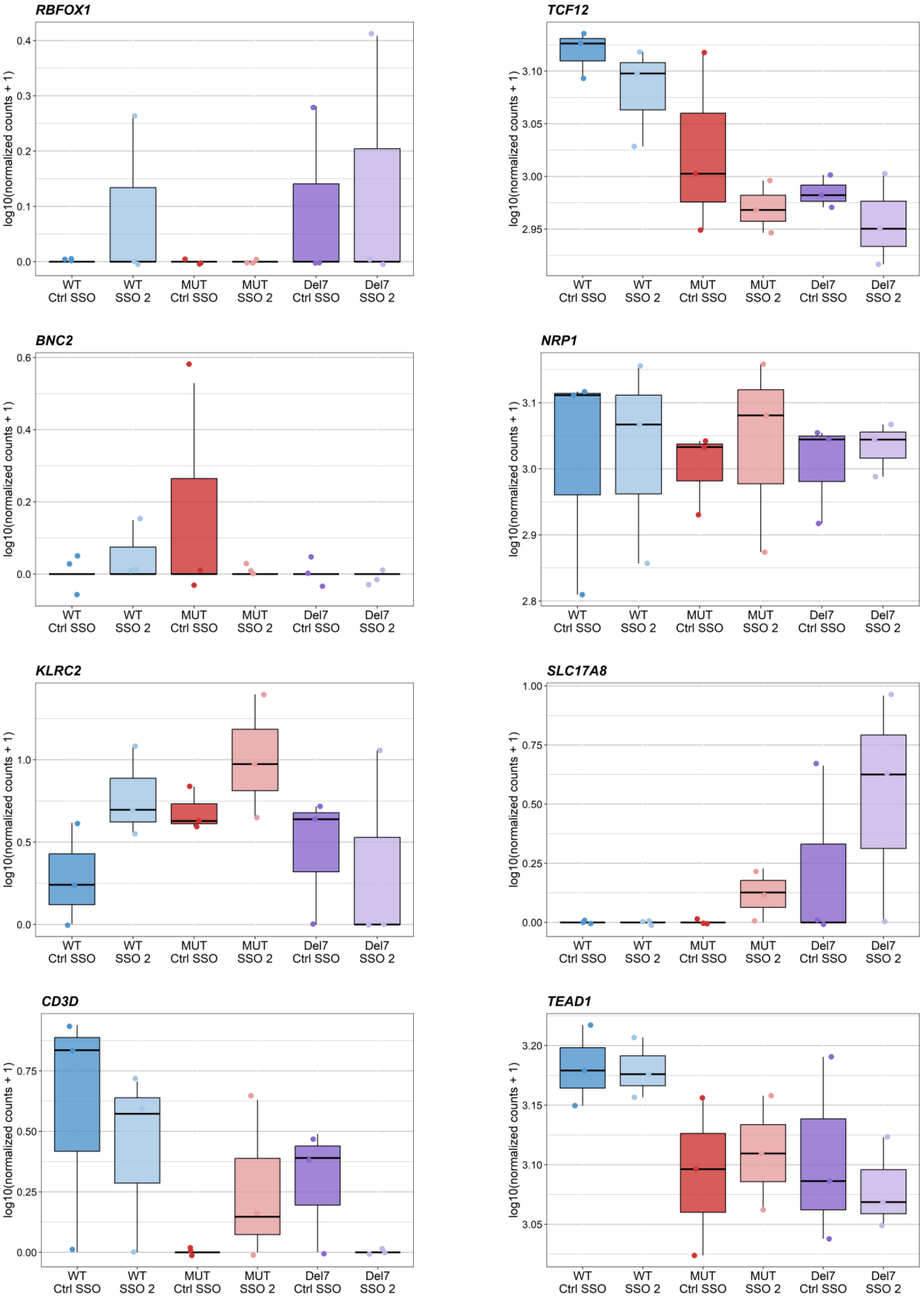

RBFOX1
TCF12
BNC2
NRP1
KLRC2
SLC17A8
CD3D
TEAD1
log10(normalized counts + 1)
WT Ctrl SSO
WT SSO 2
MUT Ctrl SSO
MUT SSO 2
Del7 Ctrl SSO
Del7 SSO 2

**Figure S11** Investigating the effect of SSO treatment on possible off-targets from control and SSO-treated CRISPR-edited HepG2 cells. Relative expression of genes with possible off-targets of SSO 2 (Table S1) between control- and SSO-treated (ctrl SSO and SSO 2) HepG2 *PCCA* WT, MUT, and Del7 cell lines indicated as log-transformed normalized read counts from RNA sequencing data (*DNAAF11* is not expressed in the CRISPR-edited HepG2 cells). *MIPOL1* is the only gene that is differentially expressed between SSO- vs. control-treated samples (Wald test with post-hoc Benjamini-Hochberg test; adjusted p-value, p.adj.=0.039), with 819 differentially expressed genes in total (p.adj.<0.05). However, no changes in the splicing patterns can be identified as affected by the possible off-target in *MIPOL1* (Figure S12), and the effect on gene expression may thus be secondary to the targeted effect from the SSO. As the SSO also blocks inclusion of the *PCCA* pseudoexon efficiently with complementarity of just 18 consecutive nucleotides in the HepG2 *PCCA* Del7 cells, complementarity in *MIPOL1* and other possible off-targets can be reduced by shortening the length of the SSO, which has been demonstrated to significantly reduce SSO off-target activity.[10] Additionally, off-target activity can be reduced by introducing mismatches to the SSO sequence, by mixing the SSO chemistry, by combining SSOs targeting the same splicing event, and by SSO delivery through free cellular uptake instead of transfection, which is also more therapeutically relevant.[10] As the SSOs used in experiments are complementary to the wild type sequence in mutant CRISPR-edited HepG2 cells and complementary to the mutant sequence in control fibroblasts, SSOs with single mismatches can efficiently block the *PCCA* pseudoexon.

### *MIPOL1*

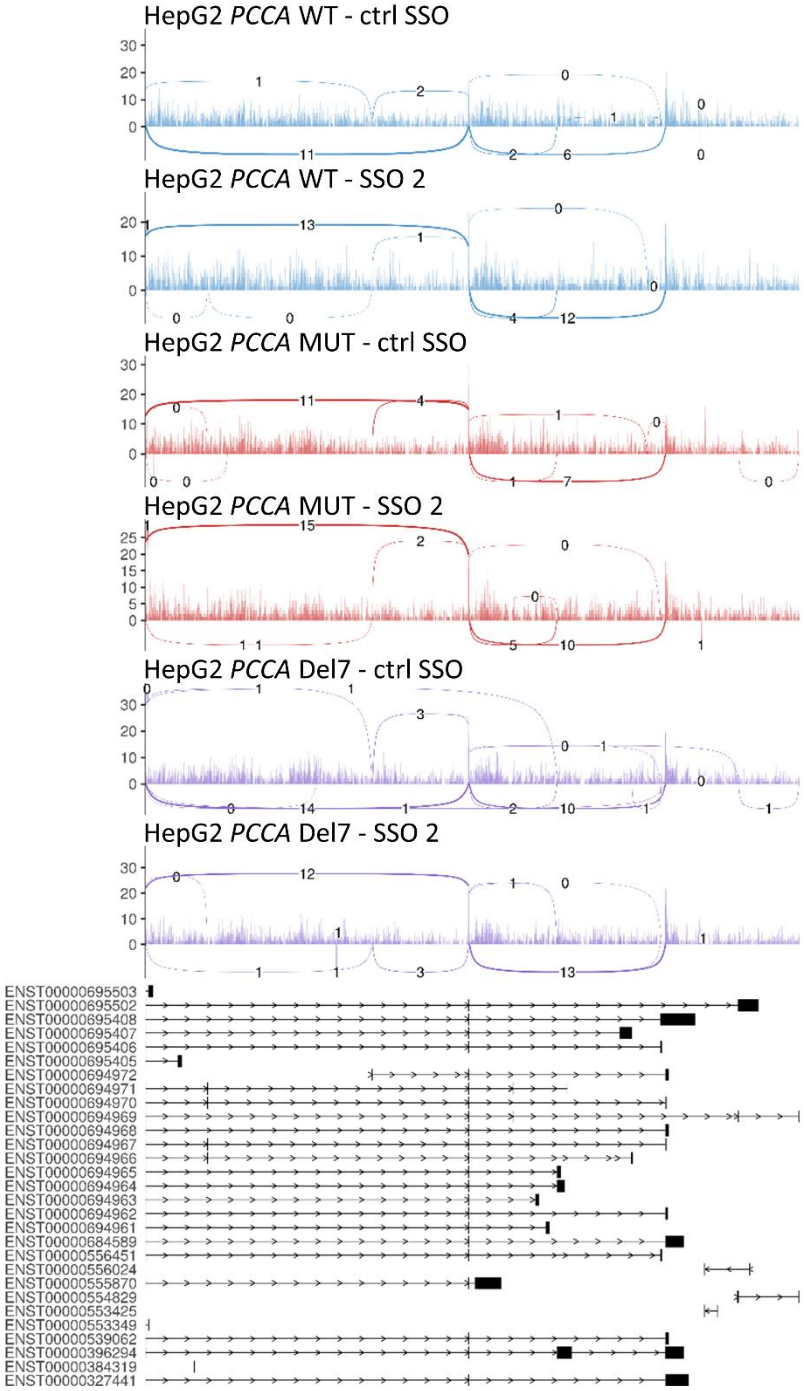


**Figure S12** Investigating the effect of SSO treatment on possible off-targets from control and SSO-treated CRISPR-edited HepG2 cells. Sashimi plots of the region from *MIPOL1* (NM_001388067; Ensemble transcript ID: ENST00000684589) exon 11 to exon 13 with aggregated read coverage (y-axis) of RNA sequencing data from control- and SSO-treated (ctrl SSO and SSO 2) HepG2 *PCCA* WT, MUT, and Del7 cell lines generated using ggsashimi (https://github.com/guigolab/ggsas). The possible SSO 2 off-target in *MIPOL1* is located in the last intron (intron 12) of the RefSeq sequence (MANE select), but is located in the last or second last exon of other transcripts (Ensembl transcript IDs, release 106: ENST00000694964 and ENST00000396294, respectively). No changes in the splicing patterns can be identified as affected by the possible off-target, and the effect on gene expression may thus be secondary to the targeted effect from the SSO.

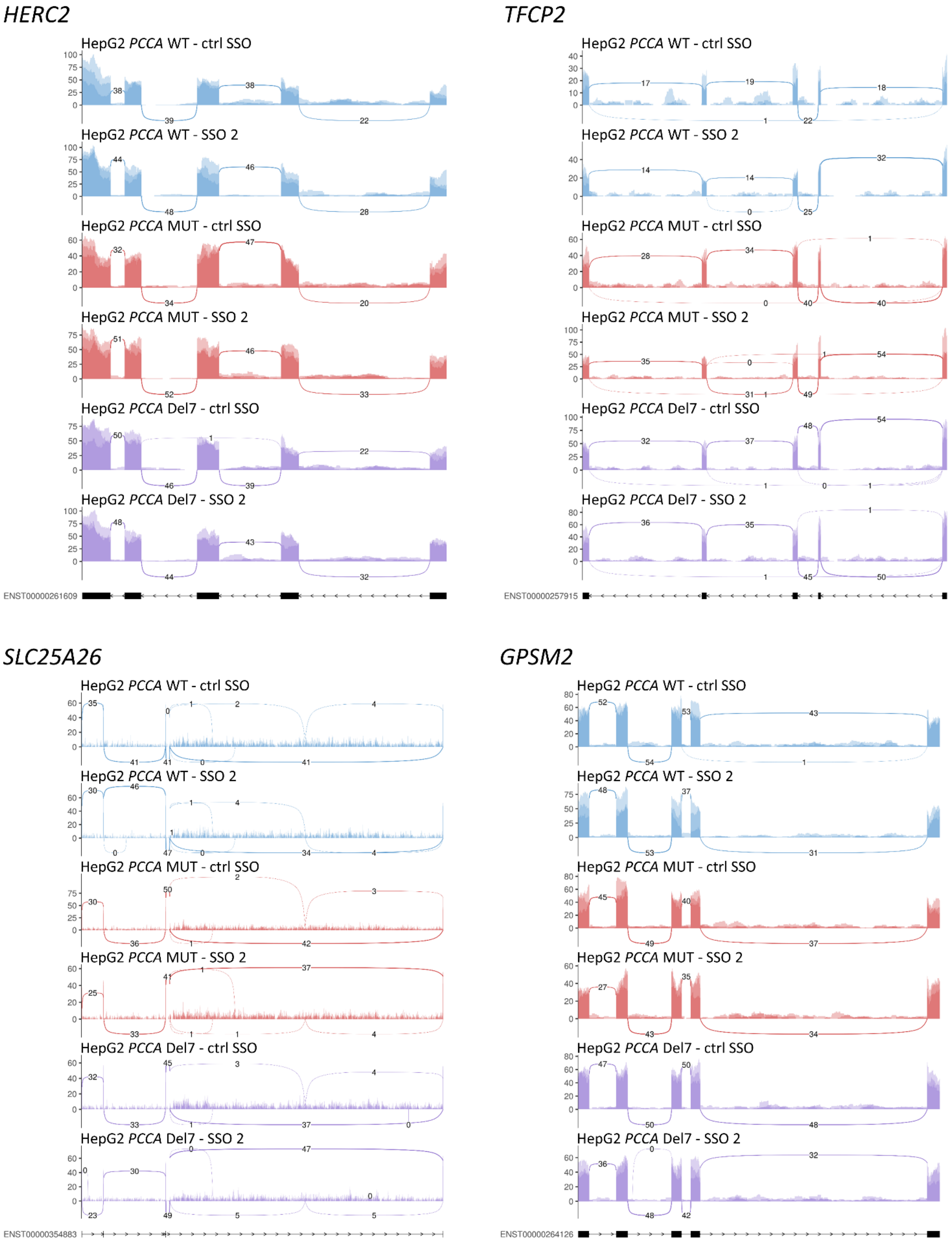
HERC2
HepG2 PCCA WT - ctrl SSO
HepG2 PCCA WT - SSO 2
HepG2 PCCA MUT - ctrl SSO
HepG2 PCCA MUT - SSO 2
HepG2 PCCA Del7 - ctrl SSO
HepG2 PCCA Del7 - SSO 2
ENST00000261609
TFCP2
HepG2 PCCA WT - ctrl SSO
HepG2 PCCA WT - SSO 2
HepG2 PCCA MUT - ctrl SSO
HepG2 PCCA MUT - SSO 2
HepG2 PCCA Del7 - ctrl SSO
HepG2 PCCA Del7 - SSO 2
ENST00000257915
SLC25A26
HepG2 PCCA WT - ctrl SSO
HepG2 PCCA WT - SSO 2
HepG2 PCCA MUT - ctrl SSO
HepG2 PCCA MUT - SSO 2
HepG2 PCCA Del7 - ctrl SSO
HepG2 PCCA Del7 - SSO 2
ENST00000354883
GPSM2
HepG2 PCCA WT - ctrl SSO
HepG2 PCCA WT - SSO 2
HepG2 PCCA MUT - ctrl SSO
HepG2 PCCA MUT - SSO 2
HepG2 PCCA Del7 - ctrl SSO
HepG2 PCCA Del7 - SSO 2
ENST00000264126

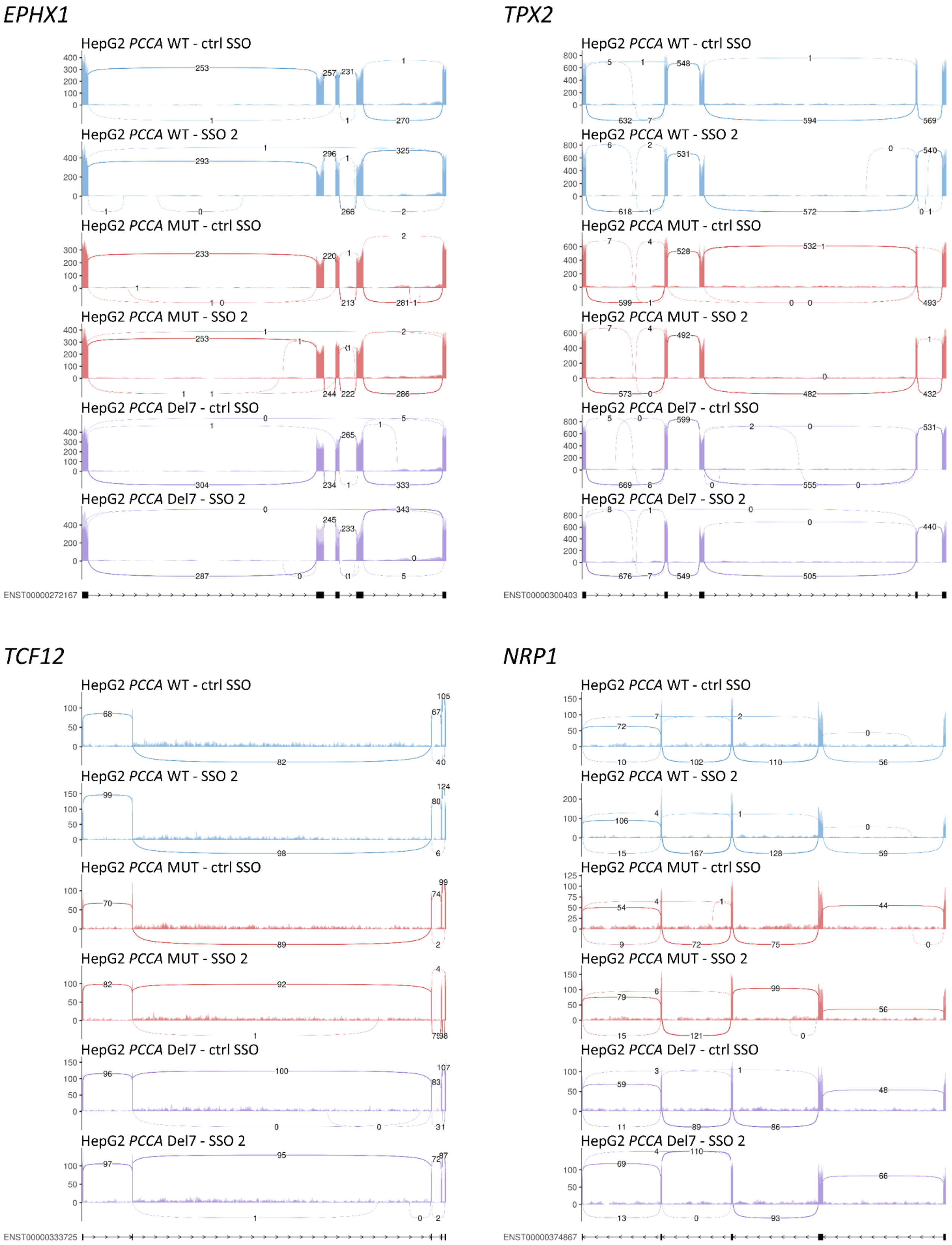
EPHX1
TPX2
TCF12
NRP1
HepG2 PCCA WT - ctrl SSO
HepG2 PCCA WT - SSO 2
HepG2 PCCA MUT - ctrl SSO
HepG2 PCCA MUT - SSO 2
HepG2 PCCA Del7 - ctrl SSO
HepG2 PCCA Del7 - SSO 2
ENST00000272167
ENST00000300403
ENST00000333725
ENST00000374867

*KLRC2*

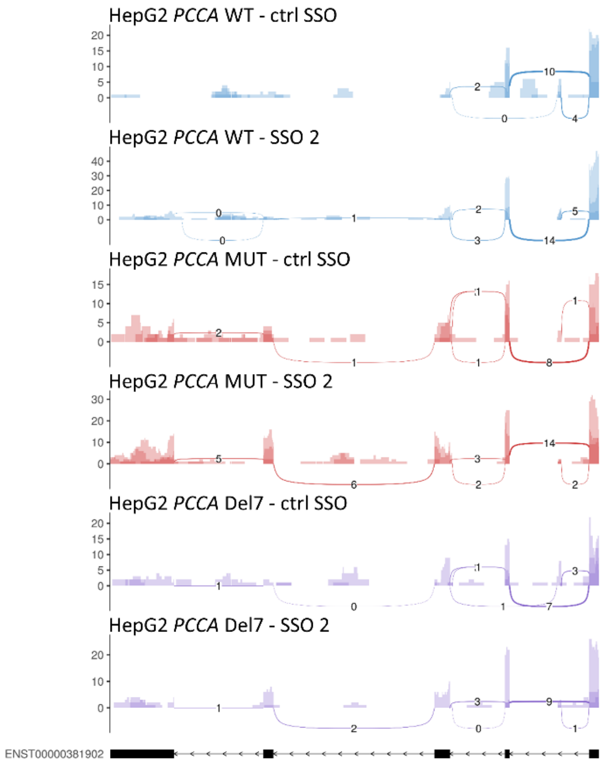


**Figure S13** Investigating the effect of SSO treatment on possible off-targets from control and SSO-treated CRISPR-edited HepG2 cells. Sashimi plots of the region from and to the second exon up- and downstream from possible off-targets of SSO 2 located in exons or <500 bp from exons in the relative RefSeq sequences (MANE Select) of the genes (Table S1); *HERC2* (NM_004667; Ensembl transcript ID: ENST00000261609; exon 74-78), *TFCP2* (NM_005653; Ensembl transcript ID: ENST00000257915; exon 8-12), *SLC25A26* (NM_001379210; Ensembl transcript ID: ENST00000354883; exon 2-6), *GPSM2* (NM_013296; Ensembl transcript ID: ENST00000264126; exon 4-8), *EPHX1* (NM_001136018; Ensembl transcript ID: ENST00000272167; exon 3-7), *TPX2* (NM_012112; Ensembl transcript ID: ENST00000300403; exon 10-14), *TCF12* (NM_207037; Ensemble transcript ID: ENST00000333725; exon 7-11), *NRP1* (NM_003873; Ensembl transcript ID: ENST00000374867; exon 8-12), and *KLRC2* (NM_002260; Ensemble transcript ID: ENST00000381902; exon 2-6). The sashimi plots are generated with aggregated read coverage (y-axis) of RNA sequencing data from control- and SSO-treated (ctrl SSO and SSO 2) HepG2 *PCCA* WT, MUT, and Del7 cell lines using ggsashimi (https://github.com/guigolab/ggsas). No changes in splicing patterns can be observed from these possible off-targets located within or near exons.

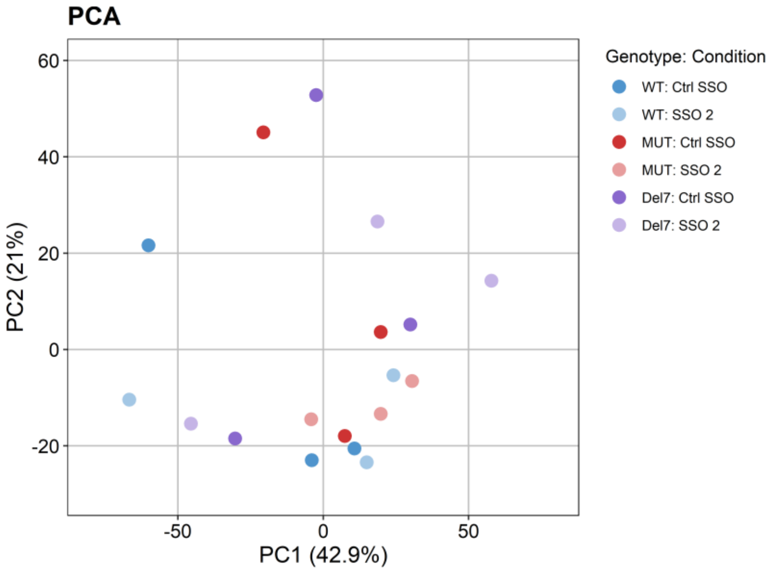


**Figure S14** Principal component analysis (PCA) plot of the variation between control- and SSO-treated (ctrl SSO and SSO 2) HepG2 *PCCA* WT (WT), HepG2 *PCCA* MUT (MUT), and HepG2 *PCCA* Del7 (Del7) cell lines in metabolomics data (n=3 from individual experiments).
